# Adipokines set neural tone by regulating synapse number

**DOI:** 10.1101/781526

**Authors:** Ava E. Brent, Akhila Rajan

## Abstract

Energy sensing neural circuits decide to expend or conserve resources by integrating tonic steady-state energy store information with phasic signals for hunger and food intake. Tonic signals, in the form of adipose tissue-derived adipokines, set the baseline level of energy-sensing neuron activity, providing context for interpretation of phasic messages. However, the mechanism by which tonic adipokine information establishes baseline neuronal function is unclear. Here we show that Upd2, a *Drosophila* Leptin ortholog, regulates actin-based synapse reorganization by reducing inhibitory synaptic contacts, thereby providing a permissive neural tone for insulin release under conditions of nutrient surplus. Unexpectedly, Insulin acts on the same upstream inhibitory neurons to conversely increase synapse number, hence re-instating negative tone. Our results suggest that two surplus-sensing hormonal systems, Leptin/Upd2 and Insulin, converge on a neuronal circuit with opposing outcomes that establish tonic, energy-store-dependent neuron activity.

**Highlights:**

- The adipokine Upd2 regulates number of inhibitory synaptic contacts on Insulin neurons.
- Upd2 activates an actin-regulating complex of Arouser, Basigin, and Gelsolin in target neurons.
- Arouser, Basigin, and Gelsolin reduce the extent of inhibitory contact on Insulin neurons.
- Insulin resets negative tone by increasing the number of synaptic contacts made by its own upstream inhibitory neurons.

## Introduction

An organism’s ability to sense different nutrient states and respond accordingly is essential to survival: when energy reserves are replete, energetically costly processes like reproduction and immunity can be invested in; under conditions of scarcity, depleted reserves signal metabolic conservation. Such homeostatic behavioral decisions are directed by a population of central nervous system (CNS), energy-store responsive neurons that receive and interpret information on systemic, steady-state energy availability from circulating adipokines. Derived from adipose tissue and released in proportion to stored fat, adipokines determine baseline target neuron activity, thereby providing the context within which the neurons integrate, interpret, and flexibly respond to additional phasic signals arising from the cyclic nutrient flux of meal intake and fasting. How energy store responsive neurons accomplish this fine-tuned integration of phasic and tonic signals is not well understood, but dysregulation of the circuits not only causes energy imbalance, but also underlies chronic metabolic disorders such as diabetes and obesity. In this study, we employ a *Drosophila* model of surplus-sensing hormones to elucidate the mechanism by which the energy-sensing circuits interpret tonic energy store status to establish baseline activity.

In vertebrates, the hormones Insulin and Leptin work together to control body weight and energy homeostasis. The circulating levels of both hormones directly correlate with total body fat mass: in conditions of energy depletion, levels decrease; in states of energy surplus, they increase (Flier, 2019; Timper and Bruning, 2017). Insulin release from the pancreatic beta cells regulates carbohydrate and lipid metabolism in peripheral tissues, promoting absorption of nutrients, such as glucose and lipids, as well as storage for their later use in the form of glycogen and triacylglycerol (TAG) (Banks et al., 2012; Flier, 2019). Concurrently, the adipokine Leptin, produced by the adipose tissue in proportion to stored fat, reports energy reserve availability to the neural circuits governing behavior. Together, Leptin and Insulin inform target neurons on energy stores (Banks, 2004; Flier and Maratos-Flier, 2017). However, Leptin and Insulin also fluctuate in a phasic manner during meal intake (Boden et al., 1996; Kolaczynski et al., 1996a; Kolaczynski et al., 1996b), making the mechanism by which the two communicate tonic information to their target neurons difficult to differentiate. Further complicating analysis is the observation that Leptin and Insulin interact with one another—and do so in sometimes synergistic, sometimes antagonistic ways. In *Drosophila*, by contrast, the relationship between Leptin and Insulin is better defined.

We previously showed that the JAK/STAT ligand Unpaired-2 (Upd2), is an adipokine that is released by the adipocytes of the fat body (FB) in proportion to fat stores, and communicates information on energy availability to the brain (Rajan et al., 2017; Rajan and Perrimon, 2012). Upd2 shares a number of features with its vertebrate counterpart, Leptin: both provide a measure of adiposity; both respond to conditions of nutritional surplus or scarcity; and both undergo restricted secretion in periods of starvation in order to conserve resources. Upd2’s primary role is controlling how much Insulin is released into circulation from the fly’s Insulin-producing cells (IPCs)--a group of 14 neuroendocrine cells, homologous to mammalian pancreatic beta cells, that reside in the median neurosecretory cluster (mNSC) of the fly’s brain (Geminard et al., 2009; Rajan et al., 2017; Rajan and Perrimon, 2012). *Drosophila* Insulin-like proteins (Dilps) not only regulate nutrient uptake and utilization, but also support many aspects of complex physiology and behavior, such as reproduction, sleep, and immunity (Das and Dobens, 2015; Enell et al., 2010; Lebreton et al., 2017; Nassel et al., 2013; Rulifson et al., 2002). Coordination of current energy status with extent of Insulin release thus ensures that the fly’s resources are properly distributed.

Upd2 signaling is received not directly by the IPCs, but by a group of proximal neurons that express the Upd2 receptor, Domeless (Dome) (Rajan and Perrimon, 2012). Binding of Upd2 to Dome activates the STAT signaling pathway in a manner that reflects current nutritional state (Rajan and Perrimon, 2012). We found that the Upd2-responsive STAT neurons are GABAergic, indicating that their function in the nutrient-sensing pathway is inhibitory: Upd2 antagonizes GABA neuron activity such that the extent of inhibition is reduced (Rajan and Perrimon, 2012). This inhibitory, clamp-like control on the effect of surplus hormones is conserved in vertebrates: Leptin-associated activation of STAT in GABAergic neurons has been shown to underpin Leptin’s effect on the hypothalamic pro-opiomelanocortin (POMC) circuits (Vong et al., 2011) that inhibit feeding and promote energy expenditure, suggesting that fat store-dependent regulation of inhibitory tone is a conserved property of Upd2/Leptin signaling. But how adipokines regulate tone is unclear.

In this study, we address that question by analyzing how Upd2 regulates the extent of inhibitory tone provided by its target GABA neurons on the IPCs. Using the pre-synaptic marker Synaptotagmin (Syt) (Yoshihara and Littleton, 2002; Zhang et al., 2002) to identify the site of synaptic contact between the Upd2-responsive neurons and the IPCs, we develop an assay for tonic neuronal activity by segmenting and quantifying Syt-expressing boutons. With this assay, we determine that Upd2 establishes inhibitory tone by regulating the number of synaptic contacts made by its target neurons on the IPCs; that several proteins involved in organization of the actin-cytoskeleton play a role in this process; and that synapse number is altered to reflect the extent of energy stores. Moreover, we demonstrate that Insulin itself provides negative feedback to the system, ensuring that inhibitory tone remains in place. Our observations open a key inroad into understanding the mechanism by which systemic hormonal signals regulate tonic neural activity in order to coordinate physiological state with behavioral response.

## Results

### A population of STAT-expressing GABA neurons synapse on the IPCs to negatively regulate systemic fat storage and Insulin release

We previously defined a population of STAT-expressing GABA neurons in the pars intercerebralis (PI) region of the *Drosophila* brain that receive fat store information from the FB in the form of the adipokine, Upd2 (Rajan and Perrimon, 2012). A clearer understanding of how these neurons regulate systemic fat storage called for a specific STAT-Gal4 driver that would recapitulate our previous observations without targeting all GABA neurons. To this end, we identified a Gal4 line from the InSite collection, in which Gal4 is inserted into the STAT92E gene (Gohl et al., 2011). We expected this line to recapitulate part, if not all, of endogenous STAT expression. With the STAT-Gal4 driver, we examined expression of dsRed in the adult brain (Figure 1A). STAT was visualized in several neuron populations, including a set of 6 neurons with somas in the PI region (left panels, yellow arrow), and arborizations in the subesophageal zone (SEZ, yellow bracket). In both location and projections, these neurons resembled the Upd2 targets we previously described (Rajan and Perrimon, 2012). We also observed populations of STAT neurons with somas in a bilateral dorsal domain (blue arrows), in the olfactory bulbs (magenta arrow), and in the SEZ (green arrow) (Figure 1A); however, our focus was the STAT-expressing neurons in the brain PI region, hereafter referred to as the PI-STAT neurons. Co-staining of dsRed with an antibody to Dilp2 revealed that the somas of the PI-STAT neurons intermingle, but do not overlap, with those of the IPCs (Figure 1A, middle panels). (Note that in Fig. 1A, middle panels, the yellow signal in the merged images is the result of maximum intensity projections and does not represent overlapping expression of STAT and Insulin.) Single XY slices through the PI region show mutually exclusive STAT and IPC somas (Figure S1), with the tracts of the PI-STAT neurons and IPCs descending together from the PI region (Figure 1A, middle and right panels, in, respectively, XY and YZ planes). We previously demonstrated that the Dome/STAT signaling pathway promotes systemic energy storage (Rajan and Perrimon, 2012). To confirm that our InSite STAT-Gal4 driver included the STAT-expressing neurons that regulate energy storage, we used the driver to induce expression of RNAi transgenes directed against either *dome* or *Stat* (Figure 1B). Any potential developmental effects arising from interference with Dome/STAT signaling were avoided by restricting our knockdown to adult *Stat*-expressing cells, using a tubulin promoter-driven, temperature-sensitive Gal80 (TubGal80^ts^), and shifting to permissive temperature only after eclosion. We found, as expected, that knockdown of either *dome* or *Stat* (STAT-Gal4, TubGal80^ts^>UAS-*dome*-RNAi or STAT-Gal4, TubGal80^ts^>UAS-*Stat*-RNAi) in *Stat*-expressing cells resulted in decreased systemic TAG storage (Figure 1B), and similarly, that over-expression of either a dominant-negative (DN) or constitutively-active (CA) (STAT-Gal4, TubGal80^ts^>UAS-*Stat-CA* or STAT-Gal4, TubGal80^ts^>UAS-*Stat-DN*) form of *Stat* resulted in, respectively, decreased or increased TAG storage (Ekas et al., 2010). To target the effects of the STAT-Gal4 driver specifically to STAT-expressing neurons, we expressed the temperature-sensitive neuronal activator *TrpA1* (Hodge, 2009) in those neurons (STAT-Gal4>UAS-*TrpA1*), and observed that at permissive temperature, systemic TAG storage progressively decreased over the course of 1 to 3 days (Figure 1C)--confirming that STAT expression in neurons is directly responsible for impacts on systemic fat storage, and that the STAT neurons regulating fat storage are inhibitory; hence GABAergic. Insulin release from the IPCs plays an essential role in promoting and regulating systemic fat storage (Das and Dobens, 2015; Rulifson et al., 2002; Zhang et al., 2009). To determine if the STAT neurons participate in Insulin release from the IPCs, we induced STAT neuron activity with TrpA1 (STAT-Gal4>UAS-*TrpA1*) and examined Dilp5 expression in the IPCs, using immunohistochemistry. It has been shown that decreased Insulin secretion can be detected as an increase in Insulin protein in the IPC somas (Geminard et al., 2009; Rajan and Perrimon, 2012). Following activation of STAT neurons with *TrpA1* for 1 day (Figure 1D), we observed retention of Dilp5 in the IPCs (a similar effect on Insulin release was visible after a shorter, 3 hour activation; data not shown). Confirming that the STAT neurons regulating Insulin release are GABAergic, we saw restored Insulin secretion when STAT-controlled *TrpA1* was co-expressed with Gad1-Gal80, which functions to repress Gal4 activity specifically in the GABAergic neurons (Figure 1D) (Sakai et al., 2009). To test if repression of Insulin release by the STAT neurons might explain the fat storage phenotype we observed following activation of the neurons with TrpA1 (Figure 1C), we asked if reduced TAG could be rescued simultaneously with induced Insulin release. Using STAT-Gal4 as well as a LexA-LexAop binary system under control of Dilp2 regulatory elements (Dilp2-LexA), we generated flies expressing *TrpA1* in both the STAT neurons and the IPCs (STAT-Gal4>UAS-*TrpA1*; Dilp2-LexA>LexAop-*TrpA1*). Following a shift to permissive temperature for 1 day, we observed that activation of the STAT neurons reduced TAG storage, while IPC activation increased it. Strikingly, double activation rescued the STAT-Gal4>UAS-*TrpA1* phenotype (Figure 1E), indicating that STAT neurons affect systemic TAG levels via their regulation of insulin release. Finally, we found that repression of STAT neurons by over-expression of the potassium channel *Kir2.1* (STAT-Gal4, TubGal80^ts^>UAS-*Kir2.1*) rescued the reduced TAG phenotype of the *upd2* mutant (*upd2Δ*) flies (Figure 1F), supporting that STAT neuron regulation of fat storage functions downstream of the Upd2 released from the FB. From these results we concluded that the STAT-Gal4 driver is able to identify and manipulate a population of GABAergic, STAT-expressing neurons that function to repress Insulin release and systemic fat storage: activation of STAT signaling in these neurons by FB-derived Upd2 relieves repression such that Insulin is released and fat stores maintained.

**Figure 1.**
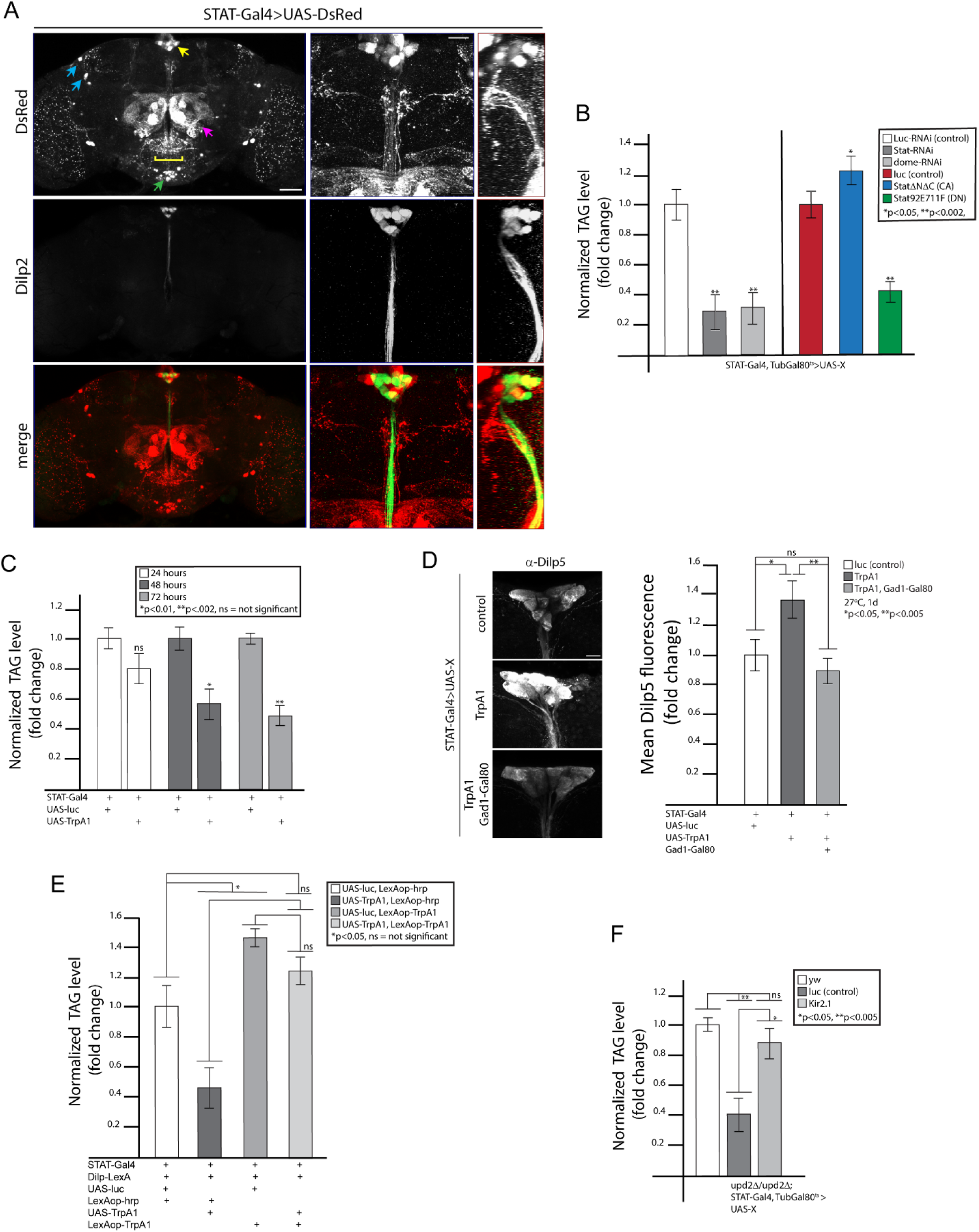
STAT is expressed in a population of PI GABA neurons that regulate systemic fat storage and Insulin release. (A) Analysis of STAT expression in the adult brain, visualized via STAT-Gal4 driving expression of DsRed. In whole brain view (left panels), STAT expression is seen in several neuron populations, including a set of 6 neurons with somas in the PI region (yellow arrow) and arborizing in the SEZ (yellow bracket), in a bilateral domain (blue arrows), in the olfactory bulbs (magenta arrow), and in the SEZ (green arrow). Intermingling of the PI STAT neurons with the IPCs is visualized with an antibody to Dilp2. At higher magnification, tracts of PI-STAT neurons and IPCs can be seen intermingling and sending tracts together from the PI region (middle and right panels, in respectively the XY and YZ planes). Scale bars: 50μm (left panels) and 20μm (middle and right panels). (B) Systemic TAG storage following manipulation of the Dome/STAT pathway in STAT-expressing neurons. (C) Systemic TAG storage following expression of *TrpA1* in the STAT neurons at 27°C for either 1, 2, or 3 days. (D) Dilp5 immunostaining in the IPCs following activation of STAT-expressing neurons with TrpA1 for 1 day at 27°C. Insulin secretion is restored when TrpA1 is repressed via Gad1-Gal80. Quantification of mean Dilp5 fluorescence indicated on the right: n=8 brains per genotype. Scale bar, 10μm. (E) Systemic TAG analysis following expression for 1 day at 27°C of either *TrpA1* in STAT neurons alone (STAT-Gal4), *TrpA1* in the IPCs alone (Dilp2-LexA), or simultaneous expression of *TrpA1* in both STAT neurons and IPCs. (F) Systemic TAG storage in *upd2Δ* mutants in which STAT neuron activity has been repressed by *Kir2.1*. CA, constitutively-active; DILP, *Drosophila* Insulin-like peptide; DN, dominant-negative; IPC, Insulin producing cell; PI, pars intercerebralis; SEZ, sub-esophageal zone; TAG, triacylglycerol.

To understand the mechanism underlying STAT-dependent disinhibition of the IPCs, our first step was locating the point of synaptic contact between the PI-STAT neurons and IPCs, using the GRASP system (GFP reconstitution across synaptic partners) (Feinberg et al., 2008; Gordon and Scott, 2009). By expressing one half of split-GFP (*spGFP*) in STAT neurons (STAT-Gal4>UAS-*spGFP1-10*, Figure 2A, first panel), and the other half in the IPCs (Dilp2-Gal4>LexAop-*spGFP11*, Figure 2A, second panel shows Dilp5 expression), we traced the reconstituted GFP to a contact point between the PI-STAT neurons and IPC tracts, just below the somas in the PI region (Figure 2A, panels 3 and 4, arrows). For visualization of the presynaptic STAT neurons in this area, we expressed a presynaptic marker, GFP-tagged *Synaptotagmin* (*Syt-GFP*), in the STAT neurons (STAT-Gal4>UAS-*Syt-GFP*) (Yoshihara and Littleton, 2002; Zhang et al., 2002), and detected Syt-GFP in a domain running along the IPC tracts--corresponding to the GRASP location we had previously observed (Figure 2B, arrows). Syt-GFP also marks the PI-STAT processes in the SEZ (Figure 2B), as well as the neurons of the olfactory bulb. A side view of the PI-STAT presynapses reveals a point of contact with the Dilp-expressing processes that project from the IPC tracts (Figure 2C). This region has previously been described as an IPC dendrite domain (Nassel et al., 2013), an observation we confirmed by examining expression of the dendrite marker DenMark in the IPCs (Dilp2-Gal4>UAS-*DenMark*) (Figure 2D) (Nicolai et al., 2010). Syt-GFP expression in the PI-STAT neurons was eliminated by Gad1-Gal80, confirming that these neurons are GABAergic (Figure 2E).

**Figure 2.**
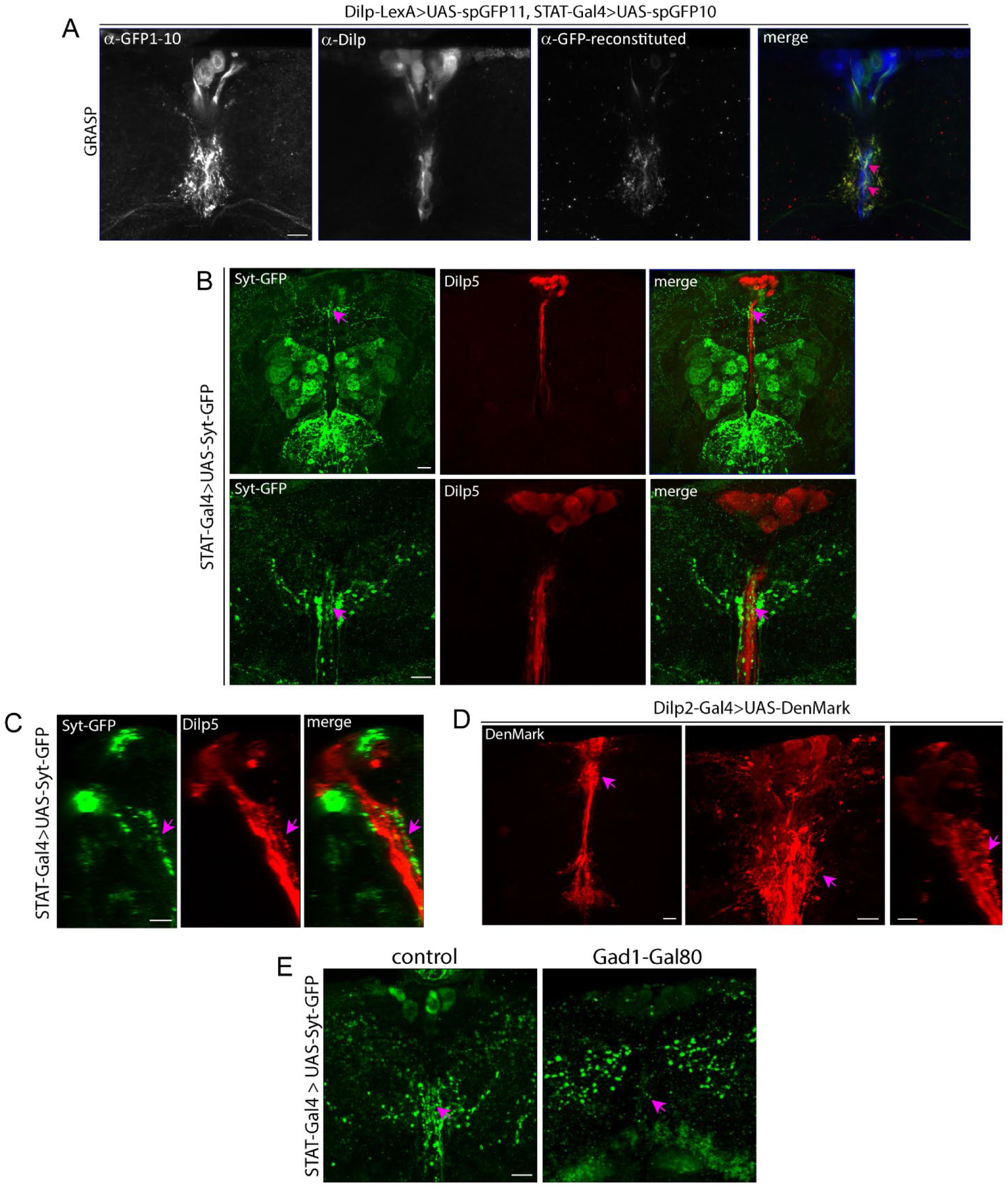
PI-STAT neurons are in synaptic contact with the IPCs. (A) GRASP detection in the PI region of adult flies with genotype STAT-Gal4>UAS-*spGFP1-10*; Dilp2-LexA>LexAop-*spGFP11*. STAT neurons identified with α-GFP antibody that specifically recognizes spGFP1-10 (first panel); IPCs identified with immunohistochemistry for Dilp5 (second panel). Points of contact (arrows, merged image) revealed by α-GFP specific to reconstituted GFP (third panel). Scale bar, 10μm. (B) Expression of Syt-GFP via STAT-Gal4 shows STAT neuron pre-synapses in adult brain. At low magnification (upper panels), Syt-GFP is visualized in the PI-STAT neurons surrounding the Dilp5-labelled IPC tracts (arrow), and in the SEZ. Expression is also seen in the olfactory bulbs. At higher magnification (bottom panels), Syt-GFP marks the PI-STAT neuron pre-synapses in contact with the IPCs (arrow). Scale bars, 20μm (upper panels) and 10μm (lower panels). (C) Side view of Syt-GFP expression in PI-STAT neurons (YZ plane) shows contact with Dilp5-expressing projections arising from the IPC tracts (middle panel). Scale bar, 10μm. (D) Visualization of Dilp2-Gal4 driving expression of UAS-DenMark in IPCs at low (left) or high (middle, XY; right, YZ) magnification identifies dendrites at point of contact between PI-STAT neurons and IPCs. Scale bars, 20μm (left panel) and 10μm (middle and right panels). (E) STAT-Gal4-driven expression of Syt-GFP in the PI-STAT neurons is eliminated by Gad1-Gal80 (arrow). Scale bar, 10μm. Dilp, *Drosophila* Insulin-like peptide, GRASP, GFP-reconstitution across synaptic partners; IPCs, Insulin producing cells; PI, pars intercerebralis; Syt-GFP, synaptotagmin-GFP.

### Establishment of presynaptic bouton number as a measurement of tonic neuronal activity

Having located the presynaptic contacts of the STAT neurons on the IPCs, we sought to analyze the mechanism by which Upd2-induced STAT signaling disinhibits Insulin release. This called for development of an assay for PI-STAT neuron activity. FB-derived Upd2 is a steady-state tonic signal, released in proportion to fat stores and continuously received by the PI-STAT neurons, which use the information to regulate repression of Insulin release in conjunction with fat stores. Thus, while the neurons are active to varying degrees depending on Upd2 secretion, they behave as a rheostat, responding to a tonic signal and steadily imparting appropriate inhibitory tone to the IPCs. As we considered tools for effective visualization of tonic activity, we reasoned that changes in baseline tone might reveal themselves as changes in the structure of synaptic contacts. It has previously been observed that alterations in synapse number or size can reflect synaptic strength (Bushey et al., 2011; Sigrist et al., 2003). We thus determined that a presynaptic marker, such as Syt, could be employed to indicate the number or size of synaptic contacts made by the PI-STAT neurons on the IPC dendrites. Syt-GFP has been used to mark bouton number in other homeostatically regulated neurons (Bushey et al., 2011; Eddison et al., 2011). Our examination of Syt-GFP expression in the PI-STAT neurons revealed Syt-GFP labeling of regularly spaced structures, likely representing boutons (Figure 3A, rectangle in left panel, middle panel). We therefore developed an image segmentation recipe to outline and analyze each Syt-GFP puncta within a region of interest, defined as the PI-STAT puncta in contact with Dilp5-labelled IPC tracts (Figure 3A-also see methods). Analysis of Syt-GFP puncta in a single brain yielded a value for average puncta number, surface area, and volume (Figure 3B). In support of this assay, we confirmed that an increase in synaptic contacts can be visualized as a greater number of Syt-GFP puncta: we examined Syt-GFP expression in PI-STAT neurons in which *EndophilinA*, an endocytic gene previously shown to cause synaptic overgrowth, had been knocked down (STAT-Gal4>UAS-*endoA*-RNAi), and observed a corresponding increase in Syt-GFP puncta (Figure S2) (Goel et al., 2019).

**Figure 3.**
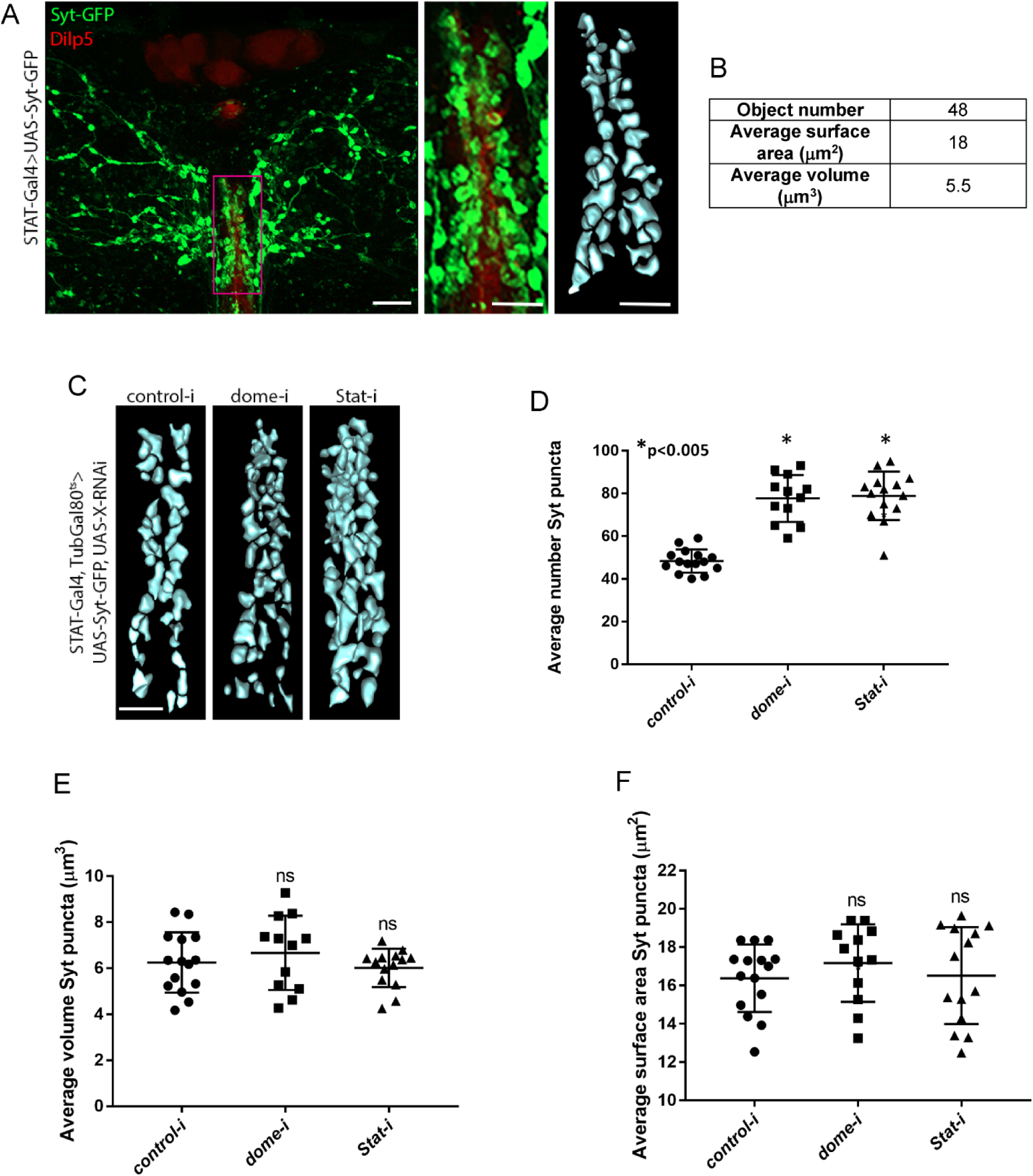
Assessment of tonic neuronal activity by segmentation analysis of presynaptic bouton number. (A) STAT-Gal4-driven expression of Syt-GFP in PI-STAT neurons. IPCs marked by immunohistochemistry with Dilp5. Box in left panel indicates region of contact between PI-STAT neurons and IPCs--also seen in middle panel. Right panel shows result of segmentation analysis. Scale bars, 10μm (left) and 5μm (middle and right). (B) Segmentation analysis of PI-STAT neuron boutons in (A). (C) Segmentation of PI-STAT Syt-GFP boutons following knock-down with *luciferase* control-, *dome*-, or *Stat*-RNAi, via STAT-Gal4, TubGal80^ts^. Scale bar, 5μm. (D-F) Average number (D), volume (E), and surface area (F) of segmented PI-STAT Syt-GFP boutons following knock-down with *luciferase* control-, *dome*-, or *Stat*-RNAi via STAT-Gal4, TubGal80^ts^. Each point represents average bouton number, volume, or surface area from a single brain. Totals of brains analyzed: for *luc*-RNAi, 15 brains; for *dome*-RNAi, 12 brains; and for *Stat*-RNAi, 14 brains. Dilp, *Drosophila* Insulin-like peptide, IPCs, Insulin producing cells; PI, pars intercerebralis; Syt-GFP, synaptotagmin-GFP.

We ascertained the suitability of Syt-GFP segmentation as a proxy for tonic neuron activity by testing what the effect would be on the Syt-GFP expressing boutons if the PI-STAT neurons were more active. We performed segmentation analysis of Syt-GFP puncta in adult brains in which either *dome* or *Stat* had been knocked down (STAT-Gal4, TubGal80ts>UAS-*dome*-RNAi or *Stat*-RNAi), and observed that in both conditions, the number of boutons increased compared to control-RNAi (Figure 3C). Examination of the segmented puncta revealed that while PI-STAT bouton number was significantly different in the absence of the Dome/STAT pathway (Figure 3D), puncta volume and surface area remained unchanged (Figure 3E-F). Altogether, our segmentation analysis suggested that the Upd2/Dome/STAT pathway regulates IPC activity by altering the number of synaptic contacts, and that quantification of bouton number would therefore provide a read-out of tonic neuron activity.

### A screen for candidate STAT target genes functioning in the PI-STAT neurons points to a role for *arouser* in the regulation of bouton number

With an assay for assessing tonic PI-STAT neuron activity established, we performed a candidate-based transgenic RNAi screen to locate genes involved in mediating STAT’s effect on bouton number. We identified a set of STAT target genes in the modENCODE consortium’s chromatin immuno-precipitation (ChIP) dataset, which was generated from a well-characterized STAT antibody in *Drosophila* embryos (Celniker et al., 2009). From the dataset, we located potential target genes with upstream STAT binding sites, and used FlyAtlas to select those expressed in the adult fly head (Table S1) (Chintapalli et al., 2007). We produced a list of 35 genes for which multiple independent RNAi lines are publicly available, and tested, via knock-down in the STAT neurons of adult flies (STAT-Gal4, TubGal80^ts^>UAS-RNAi) followed by screening for effects on systemic fat storage, which of the 35 might function in the PI-STAT neuron-IPC circuit (Table S1). Ten lines showed robust fat level changes when their activity was reduced in STAT-expressing cells, but not in adipocytes (Table S2). In a number of cases, fat storage was decreased, suggesting that these genes are positively regulated in response to Upd2 signaling. We noted that several of our identified candidates function in other physiological processes--for example, alcohol sensing, learning, and memory--all of which involve regulation of neuronal activity in response to changing internal or external stimuli. One such gene, *arouser* (*aru*), has been previously shown to participate in degree of ethanol sensitivity through modulation of bouton number (Eddison et al., 2011). Given the parallel importance of bouton number to regulation of the fat-sensing PI-STAT-IPC circuit by Upd2/Dome/STAT signaling (Figure 3C, D), we wondered if Aru might play a downstream role in the PI-STAT-IPC circuit as well.

We confirmed *aru*’s role by testing the effect of *aru* loss on fat storage. We observed reduced systemic TAG storage in three loss-of-function *aru* alleles compared to control flies (Figure 4A), and saw a similar TAG phenotype following expression of either of two independent *aru*-RNAi lines in adult fly STAT-expressing cells (Figure 4B, STAT-Gal4, TubGal80^ts^>UAS-*aru*-RNAi). Moreover, we found that over-expression of a myc-tagged version of *aru* in STAT-expressing cells (STAT-Gal4, TubGal80^ts^>UAS-*aru-myc*) led to increased TAG storage (Figure 4C). Taken together, these results indicate that Aru function in STAT-expressing cells is both necessary and sufficient to regulate systemic TAG storage. To verify that Aru functions downstream of STAT signaling, we co-expressed *aru-myc* with *Stat*-RNAi in the PI-STAT neurons of adults (STAT-Gal4, TubGal80^ts^>UAS-*aru-myc*, UAS-*Stat*-RNAi), and measured TAG levels (Figure 4D). While knock-down of *Stat* alone reduced systemic TAG, simultaneous over-expression of *aru-myc* increased systemic TAG, similar to the result we obtained when *aru* alone was over-expressed. We investigated our hypothesis that Aru’s effect on TAG storage depends on activity in STAT-expressing neurons by testing if repression of STAT neuron activity in *aru* mutants would rescue TAG storage. Following repression of STAT neuron activity with the modified Shaker K+ channel, EKO (STAT-Gal4, TubGal80ts>UAS-*EKO*) (White et al., 2001), we observed restored systemic TAG storage to WT levels (Figure 4D), indicating that Aru indeed functions in STAT-expressing neurons to control fat storage. To determine if Aru acts within PI-STAT neurons to regulate Insulin release, we examined expression of Dilp5 protein in IPCs following *aru* knockdown (STAT-Gal4, TubGal80^ts^>UAS-*aru*-RNAi), and saw significant Insulin retention (Figure 4E). Because one *aru* allele, *8-128*, results from insertion of a Gal4-containing P[GawB] element within the *aru* locus (Eddison et al., 2011), we were able to further support a role for Aru-expressing neurons in regulation of Insulin release by demonstrating that expression of dsRed under control of 8-128-Gal4 marks a population of PI neurons that resemble the PI-STAT neurons found in the same location (Figure S3). Having delineated Aru’s role in Insulin release and fat storage in STAT neurons, we speculated that Aru might be carrying out these functions through regulation of bouton number. Segmentation analysis of PI-STAT Syt-GFP puncta following *aru* knockdown (STAT-Gal4, TubGal80ts>UAS-*aru*-RNAi) showed an increase in average bouton number, while over-expression of *aru-myc* (STAT-Gal4, TubGal80^ts^>UAS-*aru-myc*) showed a decrease (Figure 4G, H). These observations indicate that Aru functions to regulate the number of PI-STAT GABAergic synapses. While we cannot rule out a role for Aru in modulating synapse activity as well, evidence against this possibility arises from our examination of a synaptophysin-tagged, pH-sensitive version of red-fluorescent Tomato (Syp-pHTomato)--a reporter of activity-dependent exocytosis (Pech et al., 2015). We expressed Syp-pHTomato in PI-STAT neurons, in combination with control- and *aru*-RNAi (STAT-Gal4, TubGal80^ts^>UAS-*aru*-RNAi), and observed that although the Syp-pHTomato domain increased after *aru*-RNAi--reflecting increased bouton number--the intensity of Syp-pHTomato fluorescence appeared qualitatively similar to that of the control-RNAi PI-STAT neurons (Figure S4). These results support that Aru functions downstream of Upd2/Dome/STAT signaling to reduce PI-STAT neuron bouton number, which in turn alters the extent to which the PI-STAT neurons provide inhibitory tone to the IPCs.

**Figure 4.**
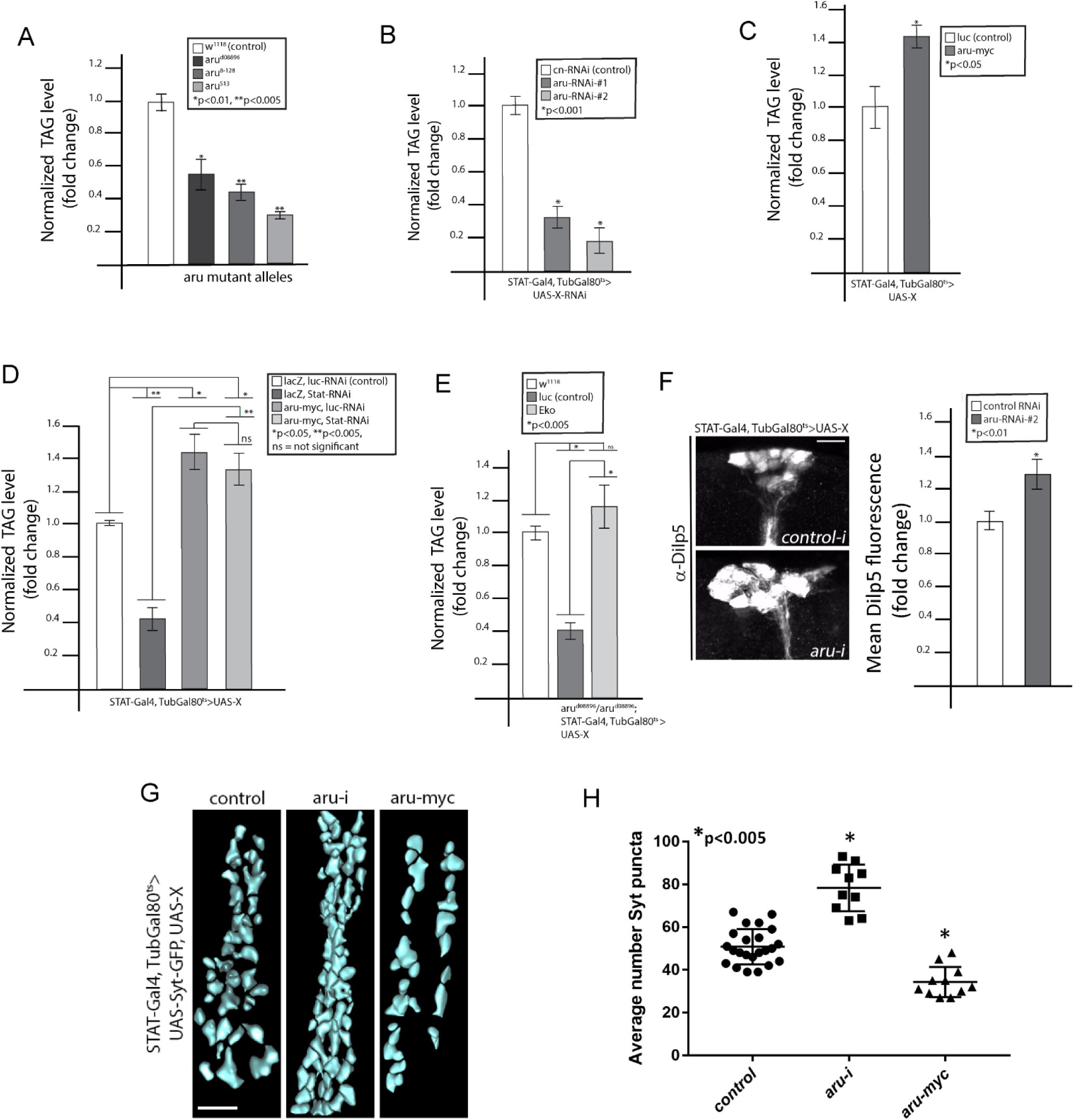
Arouser functions downstream of STAT to regulate tonic activity in the PI-STAT neurons. (A) Normalized TAG levels in control w1118 and *aru* mutant adult flies. (B) Normalized TAG levels following knockdown with control- or *aru*-RNAi in STAT neurons of adult brains, via STAT-Gal4, TubGal80^ts^. (C) Normalized TAG levels in adult flies over-expressing a myc-tagged version of *aru* in STAT neurons, via STAT-Gal4, TubGal80^ts^. (D) Normalized TAG levels following expression of both *Stat*-RNAi and *aru-myc* in STAT neurons, via STAT-Gal4, TubGal80^ts^. (E) Normalized TAG levels in aru^08896^ flies following repression of STAT neuron activity by STAT-Gal4, TubGal80^ts^-driven expression of *EKO*. (F) Immunohistochemistry for Dilp5 in IPCs of adult brains following knock-down with either control- or *aru*-RNAi in STAT neurons, via STAT-Gal4, TubGal80^ts^. Mean Dilp5 fluorescence quantified on the right: n=15 brains per genotype. Scale bar, 10μm. (G) Segmentation of Syt-GFP boutons in PI-STAT neurons following either *aru*-RNAi or *aru-myc* over-expression via STAT-Gal4, TubGal80^ts^. Scale bar, 5μm. (H) Quantification of average number of segmented PI-STAT Syt-GFP boutons in control, *aru*-RNAi and aru-myc over-expressing STAT neurons. Totals of brains analyzed: for control, 23; for *aru*-RNAi, 10; and for *aru-myc*, 11. Dilp, *Drosophila* Insulin-like peptide, IPCs, Insulin producing cells; PI, pars intercerebralis; Syt-GFP, synaptotagmin-GFP; TAG, triacylglycerol.

### Aru works together with Basigin to alter PI-STAT neuron bouton number through regulation of the actin cytoskeleton

To analyze how Aru alters bouton number in the PI-STAT neurons, we looked for the proteins with which Aru interacts, using immunoprecipitation and mass spectrometry (IP-MS). As obtaining sufficient material from brains for a robust IP-MS analysis is difficult, we performed our experiments using Aru-Myc immunoprecipitates from *Drosophila* S2R+ cells. We and others have previously demonstrated that this approach provides a relevant candidate list for verification *in vivo* (Kwon et al., 2013; Neumuller et al., 2012; Rajan et al., 2017). From our Aru IP-MS screen, we selected a set of candidates (Table S3)--among them Basigin (Bsg), an immunoglobulin domain-containing transmembrane protein that has previously been shown to participate in regulation of presynaptic cytoskeletal architecture (Besse et al., 2007). Notably, we had independently identified *Bsg* in our RNAi screen as a potential STAT-target gene: following knockdown of *Bsg* in STAT-expressing cells (STAT-Gal4, TubGal80ts>UAS-*Bsg*-RNAi), we observed decreased systemic TAG storage (Figure 5A, Table S2). To determine if Bsg, like Aru, regulates PI-STAT neuron bouton number, we performed *Bsg* knockdown in STAT-expressing cells, followed by segmentation analysis on Syt-GFP puncta (STAT-Gal4, TubGal80ts>UAS-*Bsg*-RNAi), and saw an increase in bouton number (Figure 5B and C).

**Figure 5.**
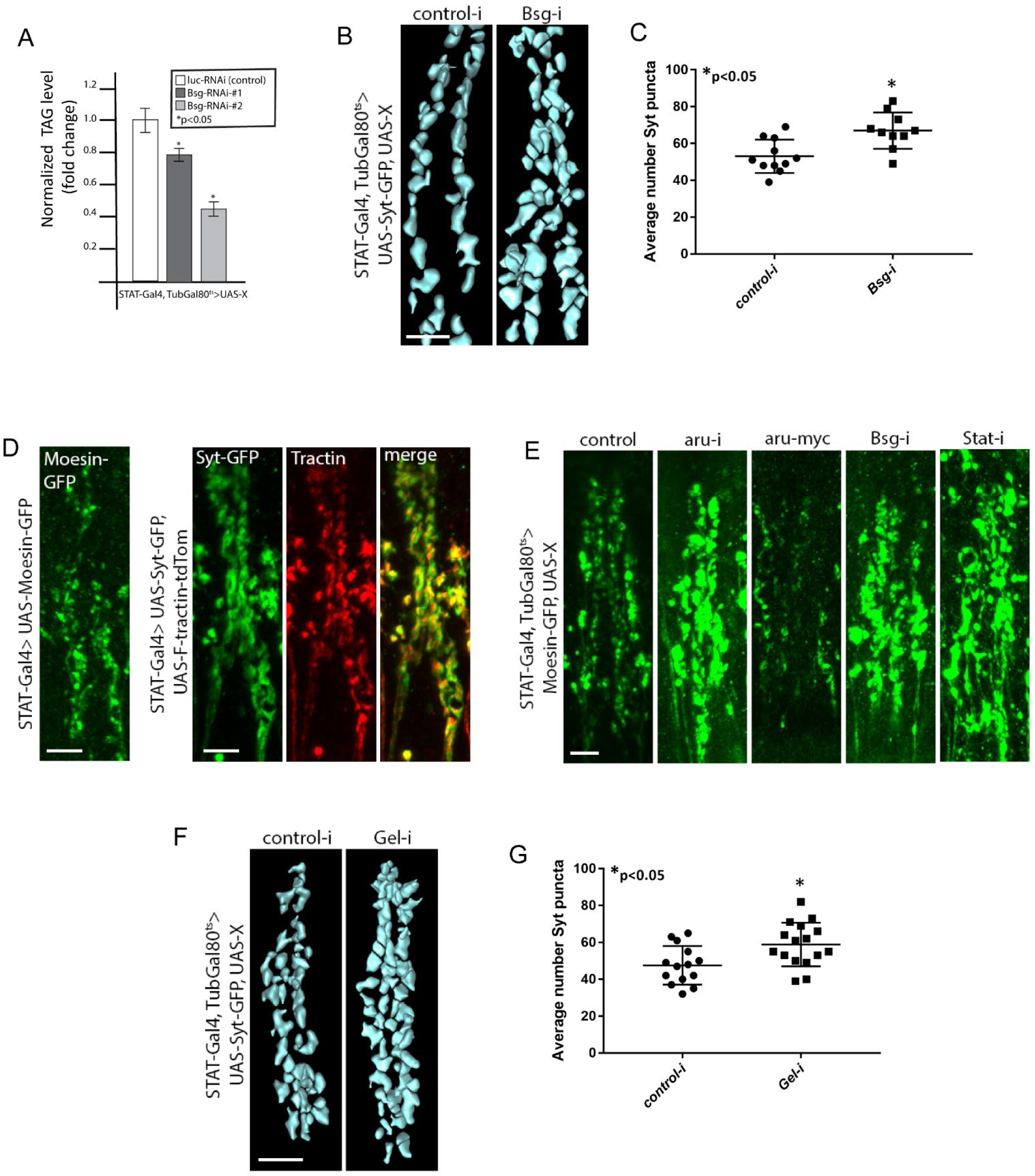
Aru and Bsg work together to regulate PI-STAT neuron bouton number through regulation of the actin cytoskeleton. (A) Normalized TAG levels in adult flies expressing either control- or *Bsg*-RNAi in STAT neurons, via STAT-Gal4, TubGal80^ts^. (B) Segmentation of Syt-GFP boutons in PI-STAT neurons expressing either control- or *Bsg*-RNAi in STAT neurons, via STAT-Gal4, TubGal80^ts^. Scale bar, 5μm. (C) Quantification of average number of PI-STAT Syt-GFP boutons following expression of control- or *Bsg*-RNAi via STAT-Gal4, TubGal80^ts^. 11 control and 10 *Bsg*-RNAi brains analyzed. (D) Left panel, Moesin-GFP expression in PI-STAT neurons, driven by STAT-Gal4. Right panels, expression of Syt-GFP and F-tractin-tdTom in boutons of PI-STAT neurons, via STAT-Gal4. Scale bar, 5μm. (E) Moesin-GFP expression in PI-STAT neurons of adult brains in which *aru*, *Bsg*, or *Stat* has been knocked-down, or *aru-myc* over-expressed, via STAT-Gal4, TubGal80^ts^. Scale bar, 5μm. (F) Segmentation analysis of Syt-GFP boutons in PI-STAT neurons following either control or *Gel*-knockdown via STAT-Gal4, TubGal80^ts^. Scale bar, 5μm. (G) Quantification of average number of PI-STAT Syt-GFP boutons in control and *Gel*-RNAi expressing STAT neurons. 14 control and 16 gel-RNAi brains analyzed. PI, pars intercerebralis; Syt-GFP, synaptotagmin-GFP; TAG, triacylglycerol.

Bsg has previously been shown to affect the presynaptic actin cytoskeleton (Besse et al., 2007). As Aru is a member of the Epidermal Growth Factor Receptor Substrate 8 (Eps8) family of proteins that also function to regulate dynamic changes in F-actin organization, we speculated that actin reorganization might underlie the bouton number phenotype we observed following manipulation of Aru or Bsg. To visualize the presynaptic actin cytoskeleton in the PI-STAT neurons, we examined expression of two genetically-tagged F-actin binding proteins, Moesin-GFP (Edwards et al., 1997) and F-Tractin-tdTomato (Tractin) (Spracklen et al., 2014), each under control of STAT-Gal4, and observed expression of both in the PI-STAT boutons (Figure 5D). Co-expression of Tractin and Syt-GFP revealed expression in an overlapping domain (Figure 5D). Following knockdown of either *aru*, *Bsg*, or *Stat*, we used Moesin-GFP to look for disruptions in the PI-STAT neuron actin cytoskeleton (STAT-Gal4, TubGal80ts>UAS-RNAi, UAS-Moesin-GFP). We observed increased and irregular expression of Moesin-GFP in the PI-STAT neurons (Figure 5E), while over-expression of *aru-myc* reduced Moesin-GFP expression (Figure 5E). These results suggested to us that Aru and Bsg regulate bouton number by altering presynaptic actin, and that Upd2 signaling triggers actin reorganization in order to reduce the extent of IPC inhibition.

Actin performs a variety of tasks in neurons, including establishment of presynaptic architecture, regulation of synaptic vesicle release, and construction of new synaptic contacts (Cingolani and Goda, 2008; Nelson et al., 2013). We asked how Aru and Bsg might alter the actin cytoskeleton to affect PI-STAT neuron activity, and hypothesized that Aru/Bsg-mediated disassembly of actin could function to eliminate presynaptic contacts, thereby reducing inhibition of the IPCs. In support of this possibility, our IP-MS experiment identified the actin-severing protein Gelsolin (Gel) as a potential Aru interactor (Table S3). A key regulator of actin filament assembly and disassembly, Gel binds to the barbed ends of actin filaments and severs existing filaments, thereby preventing monomer exchange. Because Gel’s actin-severing function has been previously shown to function during synapse elimination (Meng et al., 2015), we considered that knockdown of *Gel* in PI-STAT neurons would produce an increased bouton phenotype, similar to that observed after *aru*- or *Bsg*-RNAi. Following *Gel* knockdown (STAT-Gal4, TubGal80ts>UAS-*Gel*-RNAi), our segmentation analysis of PI-STAT Syt-GFP puncta indeed revealed an increase in average bouton number (Figure 5F and G). These results, taken together, point to a model in which FB-derived Upd2 determines the level of IPC insulin secretion by inducing the activity of an actin-regulating complex of Aru, Bsg, and Gel, which in turn functions to reduce the extent of inhibitory contact between the PI-STAT neurons and the IPCs.

### PI-STAT neuron bouton number responds to changes in nutrition and Insulin signaling

The model of an actin-based mechanism by which PI-STAT neuron bouton number is adjusted led us to ask if changes in bouton number could also be observed in response to nutrition. As Upd2 is secreted by the FB in proportion to fat stores, and increased fat stores thus yield increased Upd2 secretion, we reasoned that higher levels of circulating Upd2 should activate PI-STAT actin-remodeling to reduce bouton number and promote Insulin release. To test the theory that changes in PI-STAT bouton number reflect an increase in fat stores, we fed flies expressing *Syt-GFP* (STAT-Gal4>UAS-*Syt-GFP*) a high sugar diet (HSD), and analyzed bouton number after 1, 3, or 5 days (Figure 6A). Compared to flies fed normal food (NF), HSD fly TAG stores progressively increased over the course of 5 days (Figure 6B). Moreover, qPCR for *upd2* in the HSD fly FB tissue demonstrated, as expected, that increased fat stores resulted in higher levels of *upd2* transcription (Figure 6C). Segmentation analysis of PI-STAT Syt-GFP puncta in the brains of the flies exposed to HSD for 1, 3, or 5 days revealed a dynamic pattern (Figure 6D): After 1 day, average Syt-GFP puncta number resembled that of flies fed NF. By 3 days, average puncta number for the HSD flies was significantly lower; but by 5 days, average Syt-GFP puncta in HSD flies had returned to NF levels. These results support a model in which surplus nutrition and fat storage lead to increased levels of circulating Upd2, which in turn reduces the extent of inhibitory tone on the IPCs by decreasing PI-STAT bouton number--after which inhibitory tone is restored.

**Figure 6.**
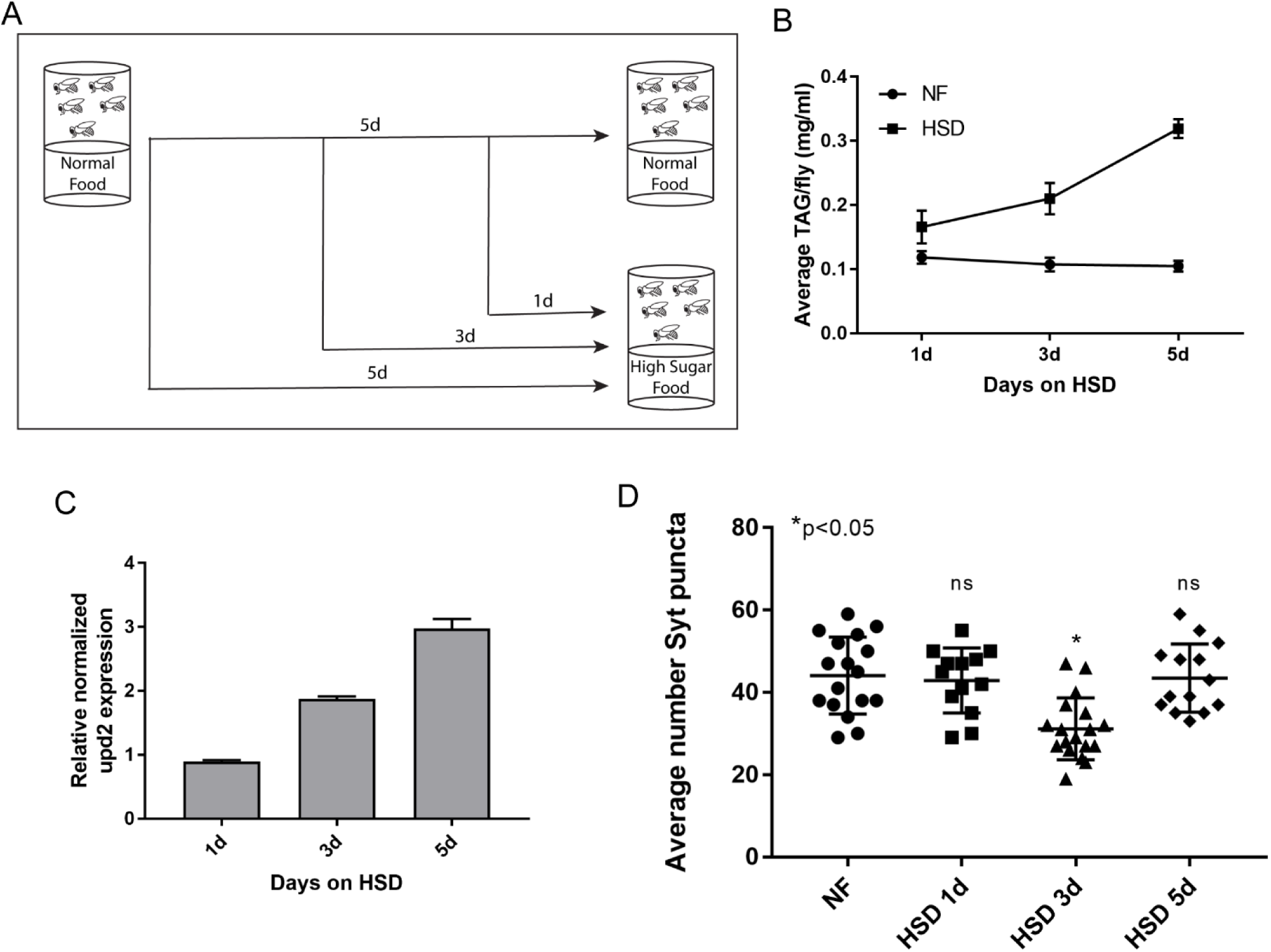
PI-STAT neuron bouton number adjusts in response to high sugar diet (HSD). (A) Time-line for HSD exposure. (B) Average TAG/fly in adult flies exposed to either NF or 1, 3, or 5 days HSD. (C) Measured by qPCR, normalized expression level of *upd2* in adult flies exposed to NF or 1, 3, or 5 days HSD. (D) Average number of Syt-GFP boutons in PI-STAT neurons of flies exposed to either NF or 1, 3, or 5 days HSD. HSD, high sugar diet; NF, normal food; TAG, triacylglycerol.

The restoration of inhibitory tone points to the possible presence of a homeostatic feedback mechanism that keeps Insulin release under negative control—a mechanism crucial to the fly’s ability to rapidly restrict Insulin secretion in conditions of nutrient deprivation. We considered that Insulin itself might be the source of the negative feedback to the PI-STAT neurons, such that rising levels of Insulin signal that inhibitory tone must be reinstated. To address this theory, we first had to determine that the Insulin signaling pathway is active in the PI-STAT neurons. From the InSite collection (Gohl et al., 2011), we identified two InR-Gal4 lines, InR-Gal4-0726 and InR-Gal4-0488, both expressed in the PI neurons as well as in the IPCs (Figure 7A and data not shown). InR-Gal4-0726 exhibits relatively restricted dsRed expression in the adult brain, in a locus primarily within the PI region in both Dilp2-expressing and non-expressing cells (Figure 7A, arrows). Examination of SytGFP in the InR-expressing cells (InR-Gal4>UAS-Syt-GFP) revealed Syt-GFP puncta resembling our previously described PI-STAT boutons in both number and location (Figure 7B, arrow in side view). Our next step was to induce Insulin signaling in adult STAT-expressing cells in order to examine the effect on systemic fat storage. In *Drosophila*, Insulin signals though a pathway that is highly conserved with that of vertebrates: Insulin binds to a receptor (InR) in target cells, thereby activating downstream targets, including phosphoinositide 3-kinase (PI3K) (Das and Dobens, 2015). Phosphorylated inositol lipids from activated PI3K serve as plasma membrane docking sites for various proteins, including a key downstream effector of Insulin signaling, AKT--resulting in activation of AKT. PI3K is additionally negatively regulated by PTEN, a PIP3 lipid phosphatase (Das and Dobens, 2015). To induce Insulin signaling in our adult STAT-expressing cells, we drove expression of activated versions of *InR* (*InR-CA*) and *PI3K* (*PI3K-CA*), and performed RNAi for *Pten* (*Pten*-RNAi) with STAT-Gal4 (STAT-Gal4, TubGal80^ts^). In all three manipulations, we observed increased systemic TAG, confirming that Insulin signaling functions in STAT-expressing cells to restrict fat storage. To identify expression of InR within the PI-STAT neurons, we made use of an antibody against human phosphorylated InR, and previously shown to cross-react with Drosophila InR (Musashe et al., 2016). We detected InR expression in the somas of GFP-marked PI-STAT neurons (STAT-Gal4>UAS-*tdGFP*) (Figure 7D, arrows), supporting a potential role for Insulin signaling in the restriction of systemic fat storage through regulation of PI-STAT neuron activity. We tested this possibility by looking at the effect of activated Insulin signaling on PI-STAT neuron bouton number. Following expression of *InR-CA* in STAT-expressing cells (STAT-Gal4, TubGal80^ts^>UAS-*InR-CA*), our segmentation analysis revealed an increase in average number of PI-STAT Syt-GFP puncta (Figure 7E and F). Consistent with this observation, activated PI3K has previously been shown to increase synapse number in the *Drosophila* neuromuscular junction (Howlett et al., 2008). Altogether, our results present strong evidence that Insulin signaling provides negative feedback to the PI-STAT neurons, thereby re-establishing the inhibitory tone that was reduced in response to changes in the level of FB-derived Upd2.

**Figure 7.**
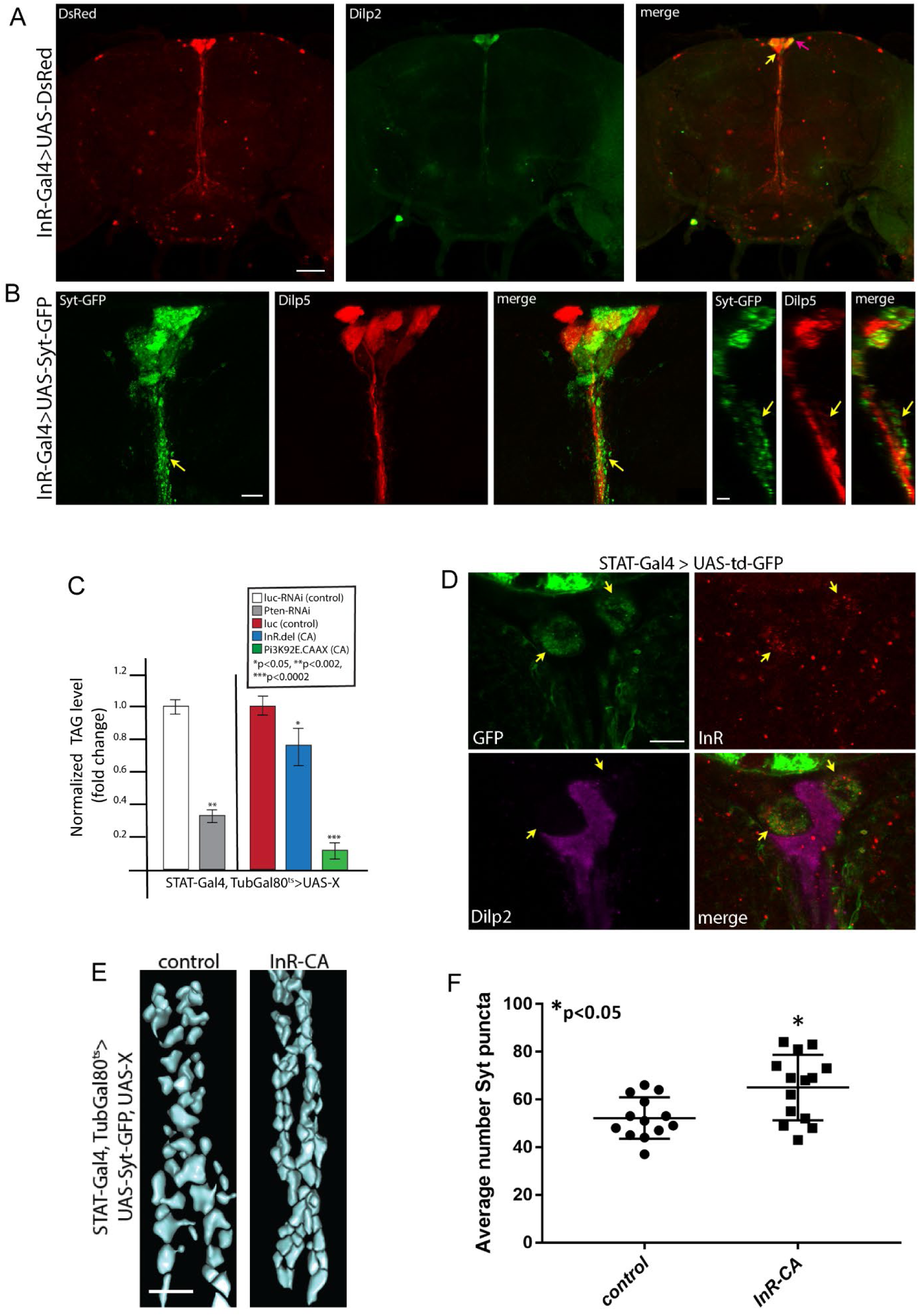
Insulin signaling re-establishes inhibitory contacts between PI-STAT neurons and IPCs. (A) InR-Gal4-driven expression of dsRed marking a population of InR-expressing neurons in the PI region. Immunohistochemistry for Dilp2 in the IPCs reveals that InR-Gal4-0726 marks both Dilp2-expressing (pink arrow) and non-Dilp2-expressing (yellow arrow) neurons. Scale bar, 50 μm. (B) Syt-GFP in InR-expressing PI neurons identifies boutons that resemble PI-STAT neuron boutons (arrows). First three panels are XY plane, last three are YZ plane. Scale bars, 10μm (XY views) and 5μm (YZ view). (C) Normalized TAG levels following knockdown of *Pten* or over-expression of constitutively active *InR* or constitutively active *Pi3K* in STAT neurons, via STAT-Gal4, TubGal80^ts^. (D) Immunohistochemistry for InR marks td-GFP-expressing PI-STAT neurons (arrows). IPCs labelled via immunohistochemistry for Dilp2. Scale bar, 10μm. (E) Segmentation analysis of Syt-GFP boutons following expression of *InR-CA* via STAT-Gal4, TubGal80^ts^. Scale bar, 5μm. (F) Quantification of average number of PI-STAT Syt-GFP boutons following expression of control or *InR-CA* via STAT-Gal4, TubGal80^ts^. Dilp, *Drosophila* Insulin-like peptide, IPCs, Insulin producing cells; InR, Insulin Receptor; PI, pars intercerebralis; Syt-GFP, synaptotagmin-GFP; TAG, triacylglycerol.

## Discussion

### Convergence of two *Drosophila* hormonal systems determines extent of inhibitory tone regulating Insulin secretion

We previously showed that the adipokine Upd2 is released from the fat body in proportion to fat stores, with the result that Insulin secretion from the IPCs in the PI region of the *Drosophila* brain is properly regulated (Rajan and Perrimon, 2012). Our observation that first-order Upd2 target neurons are GABAergic suggested a model in which STAT-expressing GABA (PI-STAT) neurons serve as a clamp on Insulin secretion, providing tonic inhibitory tone that is relieved by Upd2 signaling. In this way, Upd2, a surplus hormone reflecting fat store availability, controls another surplus hormone, Insulin, that promotes nutrient uptake and utilization as well as costly energy-expending behaviors. In the current study, we further investigate this neural circuit to address how tonic Upd2 signaling determines the extent of inhibitory tone provided to the IPCs by the PI-STAT neurons. We develop an assay to measure the number of synaptic contacts made by the PI-STAT neurons on the IPCs, and show that Upd2/Dome/STAT signaling functions to reduce the number of contacts, thereby reducing the extent of inhibitory tone on the IPCs (Figure 8A). Additionally, via a combination of genomic and proteomic screening techniques, we determine a role for reorganization of the actin cytoskeleton in the regulation of synapse number, involving Aru, Bsg, and Gel. In *C. elegans*, Gel’s actin-severing function has been shown to participate in synapse elimination under control of the cell death pathway (Meng et al., 2015). Aru’s regulation of synapse number in the *Drosophila* brain has been described (Eddison et al., 2011; LaFerriere et al., 2011); however, Aru has not heretofore been implicated in actin dynamics. And while Bsg has been previously shown to alter presynaptic actin organization such that synaptic vesicle compartmentalization and release are affected (Besse et al., 2007), our findings broaden Bsg’s known actin-dependent activities to encompass determination of synapse number. Our observations on Aru, Bsg, and Gel led us to hypothesize an energy reserve-sensing model in which STAT signaling in the PI-STAT neurons promotes expression of two target genes, Aru and Bsg, that, together with Gel, form a complex capable of regulating presynaptic actin assembly, and thereby reducing synapse number. Tonic levels of Upd2 secreted from the FB determine the extent to which the complex is active and, thus the number of synaptic contacts the PI-STAT neurons make on the IPCs. In this way, inhibitory tone reflects current energy reserves (Figure 8A-C), and in conditions of surplus, Upd2 signaling increases, driving synapse reduction and, as a result, increased Insulin secretion (Figure 8A and C).

**Figure 8.**
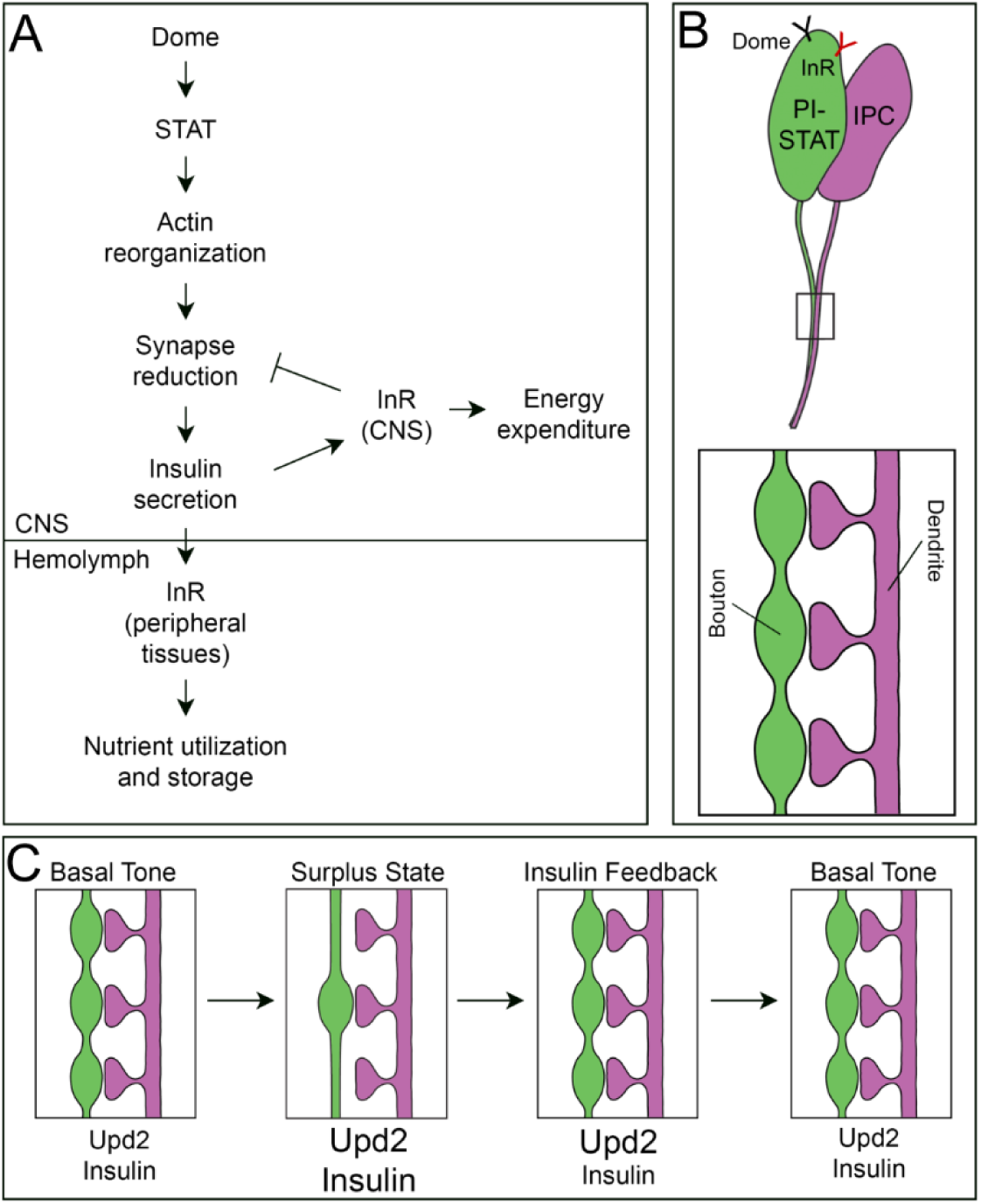
Model for convergence of two hormonal systems, Upd2 and Insulin, on the PI-STAT neuron-IPC synapse in order to regulate inhibitory tone. (A) Activation of Dome/STAT pathway in the PI-STAT neurons leads to synapse reduction through reorganization of the presynaptic neuronal actin cytoskeleton. PI-STAT synapse reduction relieves inhibition on the IPCs, leading to increased Insulin secretion. Secreted Insulin travels in the hemolymph to peripheral tissues, where it facilitates nutrient utilization and storage. Within the CNS, Insulin promotes neural regulation of energy-expending behaviors while simultaneously providing negative feedback to the PI-STAT neurons, thereby restoring inhibitory tone. (B) Somas of the PI-STAT neurons and IPCs are intermingled in the PI region of the adult brain. PI-STAT neurons express both the Dome and InR receptors. Synaptic contact occurs below the somas, in a region comprising PI-STAT neuron boutons and IPC dendrites. (C) Number of boutons in the PI-STAT neurons determines extent of basal inhibitory tone on the IPCs. Tonic levels of Upd2 set the basal tone, establishing a basal level of Insulin release. Exposure to surplus nutrition increases circulating Upd2, which reduces PI-STAT neuron bouton number and increases Insulin secretion. Elevated Insulin then feeds back on the PI-STAT neurons to re-establish tone, ensuring that Insulin secretion remains under negative control. A return to basal levels of Upd2 and Insulin occurs following a return to normal levels of nutrition. IPCs, Insulin producing cells; PI, pars intercerebralis.

Prolonged surplus nutrition and diminished inhibitory tone on the IPCs could lead to a scenario in which Insulin release is no longer under negative control; however, our finding that synapse reduction in response to surplus was subsequently reversed (Figure 6D) prompted us to search for a negative feedback mechanism that promotes PI-STAT synapse number. We found that the insulin pathway is expressed in the PI-STAT neurons, and that activation of InR by Insulin indeed promotes synapse number, which then re-establishes inhibitory tone on the IPCs (Figure 8A and B). We propose that this negative feedback mechanism ensures that increases in Insulin secretion are not long-term, and that restoration to a basal level of Upd2 is not dependent solely upon reduced food intake. Instead, Upd2 and Insulin work together in the PI-STAT-IPC circuit to generate an energy-sensing program that steadily maintains the extent of basal inhibitory tone on Insulin release, despite fluctuations in surplus nutrition intake (Figure 8c). Since high levels of Insulin secretion during a fasting/starved state would threaten survival, this dual hormone inhibitory control system is essential. Moreover, the reset of inhibitory tone by negative feedback may serve as a mechanism to prevent the peripheral Insulin resistance that can develop from high levels of circulating Insulin.

It is worth noting that while we have identified a mechanism for PI-STAT neuron regulation of the IPCs that involves reorganization of the actin cytoskeleton, it is likely that there are other mechanisms that do not involve Aru and Bsg. Identifying these additional energy-sensing processes will be an important area of future study.

### Conservation of mechanisms for adipokine function

While various aspects of energy-sensing differ between invertebrates and vertebrates, the primary physiological roles of Upd2 and Leptin demonstrate convergence. Both are adipokines that provide energy-store information to the CNS circuits regulating energy expenditure and meal intake, and in conditions of starvation, the circulating levels of both Upd2 and Leptin must be reduced to conserve energy for survival (Ahima et al., 1996; Rajan et al., 2017; Rajan and Perrimon, 2012). Moreover, while vertebrate neuronal circuitry under control of Leptin signaling is more complex than the PI-STAT-IPC circuit we have identified in flies, there are informative similarities there as well. Leptin alters the activities of the agouti related peptide (AgRP) and POMC target neurons, which act respectively to promote and reduce food intake (Timper and Bruning, 2017). Leptin inhibits the AgRP neurons and enhances the POMC neurons, establishing a system of dynamic regulation by energy status that matches feeding behavior to nutritional demands. Several studies suggest that the bulk of Leptin’s anti-obesity function is mediated by its direct effect on the GABAergic neurons that synapse on and inhibit the hypothalamic POMC neurons (Vong et al., 2011; Xu et al., 2018). These GABA neurons include the AgRPs, which inhibit POMC neuron activity and are themselves GABAergic (Xu et al., 2018). Thus, like the IPCs we have described in *Drosophila*, the POMCs are under GABAergic inhibitory tone, and inhibition is relieved by Leptin signaling (Vong et al., 2011).

Most analyses of Leptin’s effect on mammalian target neurons have focused on Leptin’s role in altering its neurons’ polarization state--hence their intrinsic excitability--via regulation of the hyperpolarizing potassium channels (Baver et al., 2014; O’Malley et al., 2005; Spanswick et al., 1997; Yang et al., 2010). It is likely that this modulation of excitability reflects Leptin’s additional mammalian function as a communicator of phasic signals—the signals that fluctuate with cycles of meal intake and fasting--and thus produce only short-term changes in target neuron function. While there are indications that Leptin might use a synaptic contact dependent mechanism to regulate target neurons (Pinto et al., 2004), similar to that we describe for Upd2, distinguishing between Leptin’s phasic and tonic effects has been difficult. In *Drosophila*, however, there is evidence to suggest that phasic and tonic reporting of nutrient flux is divided between Upd2 and a second Leptin ortholog, Upd1 (Beshel et al., 2017). Unlike Upd2 and Leptin, Upd1 is not expressed in the fat; it is found instead in the brain, where it has been described as a reporter of satiety to the NPF neurons that sense hunger and promote foraging behaviors and food intake (Beshel et al., 2017). While the peripheral signal activating release of Upd1 has not been identified, loss of the satiety signal leads to over-eating and an obese phenotype. Notably, Drosophila NPF is the equivalent of the mammalian hormone NPY, produced by the AgRP neurons to promote food intake (Beshel and Zhong, 2013; Wu et al., 2005b). These observations open the possibility that post-meal phasic information on energy availability is transmitted through Upd1, and tonic FB energy store availability through Upd2-- and that the two converge to affect Insulin release via the PI-STAT neurons or other circuits. While the relationship between Upd1 and Upd2 remains an avenue for future research, our examination of Upd2’s unique role in reporting adipose tissue energy reserves provides key mechanistic insights into adipokine-dependent regulation of neural tone.

### A role for Eps8-family members in regulation of synapse number

We have identified a role for Aru in determining synapse number in the PI-STAT-IPC circuit— paralleling Aru’s previously described regulation of synapse number with regard to ethanol sensitivity (Eddison et al., 2011). An IP-MS experiment allowed us to query the mechanism by which Aru performs this function. We identified a set of Aru-interacting proteins—including Bsg and Gel, described in this study, as well as several members of the Arp2/3 actin-nucleation complex (data not shown)--which point to Aru’s regulation of the actin cytoskeleton: examination of actin organization following manipulation of Aru and Bsg suggests that these proteins may disassemble the cytoskeleton to effect synapse elimination.

Aru is a member of the Eps8 family of predicted adaptor proteins (Eddison et al., 2011; Tocchetti et al., 2003). In mammals, this family comprises Eps8, Eps8L1, Eps8L2, and Eps8L3 (Tocchetti et al., 2003)--the best-characterized of which is Eps8, whose role in actin dynamics has been extensively studied (Disanza et al., 2004; Disanza et al., 2006; Frittoli et al., 2011; Offenhauser et al., 2004; Roffers-Agarwal et al., 2005). Eps8 family members share a modular organization consisting of several protein-binding domains as well as a C-terminal effector domain which binds actin directly. Through this domain, and together with several interacting protein partners, Eps8 performs a number of tasks that affect actin organization, including actin polymerization, capping, and bundling. *Drosophila* Aru is most closely related to mammalian Eps8L3--the only Eps8 family member lacking the actin-binding domain (Disanza et al., 2004; Disanza et al., 2006; Innocenti et al., 2003; Scita et al., 2001; Tocchetti et al., 2003). Of interest, the *Drosophila* genome contains a second Eps8 member, LP01469p/CG8907, which contains all of the Eps8 protein domains, including the actin-binding domain (Tocchetti et al., 2003). That the Eps8 family consists of members with and without the C-terminal actin-interacting domain raises the possibility that the full-length and truncated proteins serve distinct functions. In support of this idea, it is notable that *Eps8* mutations in mice cause an ethanol sensitivity phenotype opposite to that of *aru* mutant flies: while *aru* mutants exhibit hypersensitivity to ethanol exposure, the sensitivity of the *Eps8* mutants is reduced (Eddison et al., 2011; Offenhauser et al., 2006). Similarly, while we have described a lean phenotype for Drosophila *aru* mutants, GWAS studies have identified variants in human *Eps8* that correlate with abnormal amounts of circulating insulin and cardiovascular risk factors for traits that include waist measurement and fat storage (Melzer et al., 2008; Schurks et al., 2011; Simonson et al., 2011; Smith et al., 2010). These findings suggest that *Eps8* may be involved in regulation of human fat storage, and--as observed in the case of alcohol sensitivity--may function in opposition to *aru*. It is significant that both *Eps8* in mammals, and *aru* in Drosophila, participate in organization of the actin cytoskeleton; in fact, the alcohol resistance phenotype observed for *Eps8* mutants has been shown to depend on actin cytoskeleton reorganization (Offenhauser et al., 2006). In view of the role we have described for Aru in inhibiting F-actin assembly, it is possible that full-length and truncated Eps8 family members antithetically affect actin reorganization. In support of this hypothesis, we have observed that over-expression of *CG8907* in Drosophila S2R+ cells results in an increase in F-actin organization (data not shown)—in direct contrast to the phenotype we found following over-expression of *aru* in STAT neurons.

### Body weight set point, alcohol sensitivity, and the physiology of tolerance

In this study, we have identified a mechanism by which Upd2 establishes a basal level of inhibitory tone to control Insulin release through regulation of synapse number. We have additionally demonstrated that while increases in surplus nutrition can reduce inhibitory tone, negative feedback from Insulin itself re-establishes negative regulation of Insulin secretion. Homeostatic levels of energy-sensing neuron activity are thus maintained--as long as the cycles of meal intake and fasting fall within normal parameters. If animals undergo chronic exposure to surplus nutrition, however, the inhibitory PI-STAT-IPC circuit may be unable to return to former basal levels. With progressively increasing energy stores, Upd2 secretion, and demands for Insulin, restoration of inhibitory tone via Insulin-dependent negative feedback would still occur--but at a higher basal level of Insulin secretion. This hypothesis has implications for the “set-point” concept of energy homeostasis, which proposes that body weight will be maintained at a stable range despite short-term variability in energy intake and expenditure (Chapelot and Charlot, 2019). Maintenance of set-point occurs via homeostatic negative feedback processes, such as the Insulin feedback mechanism described in this study. If set-point is challenged over long periods of chronic surplus nutrition, however, it may shift such that feedback mechanisms maintain energy homeostasis at a higher set-point, as may occur in obesity.

The concept that chronic challenges to energy-sensing can alter homeostatic baseline parallels the pattern of resistance observed following chronic consumption of alcohol. In this context, it is noteworthy that regulation of synapse number by Aru has been shown to affect the response to ethanol: *aru* mutants exhibit hypersensitivity, suggesting that Aru functions in the development of tolerance (Eddison et al., 2011). Increased resistance to the effects of alcohol has additionally been shown to occur via altered neuronal actin dynamics--similar to the regulation of energy-sensing neuronal circuits described here (Offenhauser et al., 2006; Sordella and Van Aelst, 2006). These parallels pose the possibility that tolerance to chronic surplus nutrition (new set point) and alcohol consumption may develop through the same mechanism. In support of this connection, an FDA-approved drug for weight-loss, topiramate, has also been successfully employed to treat alcohol addiction (Caricilli et al., 2012; Cosentino et al., 2013; Johnson and Ait-Daoud, 2010).

### Experimental Procedures

#### Drosophila Strains

We used the *upd2* homozygous deletion mutant, *upd2D3-62* (*upd2Δ*; (Hombria et al., 2005)). The *aru^08696^*, *aru^8-128^*, and *aru^S13^* mutant strains were provided by Ulrike Heberlein (Eddison et al., 2011; LaFerriere et al., 2011). UAS and LexAop lines: UAS-*luc* (Rajan and Perrimon, 2012); UAS-*dsRed2* (BDSC #8546); UAS-*DenMark* (BDSC #33061); UAS-*TrpA1* (BDSC #26263); UAS-*EKO* (BDSC #40974); UAS-*tdGFP* (BDSC #35839); UAS-*Syt-GFP* (BDSC #6925); UAS-*Moesin-GFP* (BDSC #31775); UAS-*F-Tractin-tdTomato* (BDSC #58989); UAS-*InR-CA* (BDSC #8254); UAS-*PI3K-CA* (BDSC #8294); LexAop-*Hrp* (BDSC # 56523); UAS-*Kir2.1* (provided by Richard Baines) (Baines et al., 2001); LexAop-*TrpA1* provided by Barret D. Pfeiffer; UAS-*Synaptophysin-pHTomato* provided by Andre Fiala (Pech et al., 2015); UAS-*spGFP1-10* and LexAop-*spGFP11* provided by Kristin Scott (Gordon and Scott, 2009); UAS-*STAT92E^ΔNΔC^* (CA) and UAS-*STAT92E^Y711F^* (DN) provided by Erika Bach (Ekas et al., 2010); Gad1-Ga80 provided by Toshihiro Kitamoto; and UAS-*aru-myc* (generated for this study). Gal4 and LexA lines: STAT-Gal4 (IT.GAL4 [0092-G4]; BDSC #62634) (Gohl et al., 2011); Dilp2-GAL4 (Wu et al., 2005a); InR-Gal4 (IT.GAL4 [0726-G4]; BDSC #63762) (Gohl et al., 2011); and Dilp2-LexA (generated for this study). RNAi lines: *luciferase*-RNAi (TRiP JF01355; BDSC #31603); *Stat92E*-RNAi (TRiP JF01265; BDSC #31317); *dome*-RNAi (TRiP JF01269; BDSC #31245); *endoA*-RNAi (TRiP JF0275; BDSC #27679); *cn*-RNAi (VDRC KK101938; #V105854); *aru*-RNAi-#1 (VDRC KK109173; #V105755); *aru*-RNAi-#2 (NIG 4276R-1); *Bsg*-RNAi-#1 (VDRC GD15718; #V43306); *Bsg*-RNAi-#2 (TRiP HMC03195; BDSC #52110); *Pten*-RNAi (TRiP JF01987; BDSC # 25697); *Gel*-RNAi (TRiP JF01723; and BDSC #31205).

### Food and Temperature

Flies were cultured in a humidified incubator at 25°C on standard lab diet, containing per liter: 15 g yeast, 8.6 g soy flour, 63 g corn flour, 5g agar, 5g malt, 74 mL corn syrup. High sugar diet consisted of standard lab diet plus 30% additional sucrose by volume. Males only, age 7-15 days post-eclosion, were used in all experiments. For RNAi experiments with TubGal80^ts^, crosses were maintained at 18°C for 7 days post-eclosion, after which the progeny were shifted to 29°C for 5-7 days. For RNAi experiments without TubGal80^ts^, crosses were placed at 29°C until time of analysis. For TrpA1 experiments, crosses were maintained at 18°C until 7 days post-eclosion, after which they were transferred to 27°C for 1-3 days. For EKO and Kir2.1 experiments with TubGal80^ts^, crosses were maintained at 18°C until 7 days post-eclosion, after which they were transferred to 29°C for 3 days.

### RNAi Screen for STAT Target Genes

Candidate STAT targets were identified from a modENCODE STAT ChIP dataset (Celniker et al., 2009). Hits with a STAT binding site within 500bp of the transcriptional start were selected, and those showing expression in the adult brain on the FlyAtlas database were used for further analysis (Table S3) (Chintapalli et al., 2007). Multiple independent RNAi lines were obtained for each candidate, and knockdown was performed in STAT-expressing cells (STAT-Gal4). Effect on fat storage was examined by TAG assay. To ensure that effects on fat storage were not due to activity within the fat tissue itself, we additionally knocked down each candidate within the adipocytes, using the FB-specific driver, Lpp-Gal4 (Brankatschk and Eaton, 2010). Table S2 lists the potential STAT target genes that affected fat storage when knocked down in STAT-expressing cells, but not in adipocytes.

### aru-myc and Dilp2-LexA Cloning and Transgenic Flies

All cloning was done using the Gateway® Technology. *arouser* cDNA cloned into the entry vector (pDONR223-FlyBiORFeome-GE009432) was obtained from the FlyBiORFeome collection maintained at the DGRC. Using LR clonase reaction (Gateway® LR Clonase® II Enzyme mix, Cat#11791-020, Invitrogen), the entry vectors were moved into destination vectors compatible with fly transformation, protein production, or cell culture, and with the appropriate N-terminal tags. For Dilp2-LexA, primers 5’ – CACCGCGTGCAACTCGACAATC-3’ and 5’ – AGGTTGCTTTACGATCAAATG - 3’ were used to amplify a region upstream of Dilp2, flanking sequences GCGTGCAACTCGACAATC and AGGTTGCTTTACGATCAAATG, to generate a 2047bp product. The PCR product was cloned into pENTR-D/TOPO and transferred to vector pBPnlsLexA-GADflUw-DEST by gateway cloning to generate the Dilp2-LexA flies. Dilp-LexA was verified by crossing to LexAop-GFP reporter, and by immunohistochemistry for Dilp5 and Dilp2. Transgenic fly lines were generated by Rainbow Transgenic Flies, Inc.

### Generation of Insulin Antibodies

Anti-Dilp5 and chicken anti-Dilp2 primary antibodies were developed by New England Peptide (NEP). Peptide H_2_N-CPNGFNSMFA-OH was injected into rabbits for Dilp5, and Ac-CEEYNPVIPH-OH into chickens for Dilp2.

### Immunoprecipitation and Mass Spectrometry

For Immunoprecipitation (IP) from S2R+ cells, protein for each condition was prepared by lysing 2 wells of a 6-well dish, 4 days after transfection. 2mg/ml was used per IP experiment, performed with camelid antibodies Myc-Trap™ Magnetic Agarose (cat# ytma-10, Chromotek), following manufacturer’s protocol. The sample was prepared for mass-spec as previously described (Rajan et al., 2017). Immunoprecipitates were electrophoresed approximately one to two centimeters into a SDS-PAGE gel, and the stained gel band was cut out and proteolytically digested with trypsin as described (Cheung et al., 2017). Desalted peptides were subjected to LC-MS/MS with an OrbiTrap Elite mass spectrometer, and collected data were analyzed by Proteome Discoverer v2.2. Identified peptides were filtered to a 1% FDR.

### Triglyceride Measurements

TAG assays carried out as previously described (Rajan et al., 2017). Three adult males were used per biological replicate. Note: For adult TAG assays, the most consistent results, with lowest standard deviations, were obtained with 10 day old males. Reagents for the assay were obtained from Sigma: Free glycerol (cat # F6428-40ML), Triglyceride reagent (cat# T2449-10ML), and Glycerol standard (cat# G7793-5ML). TAG readings from whole fly lysate were normalized to number of flies per experiment. The normalized ratio from the control served as baseline, and the data is represented as fold change of experimental genotypes with respect to the control. Statistical significance was quantified by 2-tailed t-test on 3-6 biological replicates per condition. Error bars indicate %SD (Standard Deviation).

### qPCR for *upd2*

qPCR Total RNA was prepared from 12-15 fat bodies per genotype, using the Direct-zol RNA miniprep kit (Zymo Research, cat#R2071). cDNA was prepared using iScript cDNA Synthesis (Bio-Rad, cat#1708891), and 1 mg RNA was used per reaction. qPCR was performed with iQ SYBR Green Supermix (Bio-Rad, cat#1708882). *rpl13A* and *robl* were employed to normalize RNA levels. Relative quantification of mRNA levels was calculated with the comparative CT method. Primers: *rpl13A*: 5’ – AGCTGAACCTCTCGGGACAC - 3’ and 5’ – TGCCTCGGACTGCCTTGTAG - 3’; *robl*: 5’ – AGCGGTAGTGTCTGCCGTGT – 3’ and 5’ – CCAGCGTGGATTTGACCGGA - 3’; and *upd2*: 5’ - CGGAACATCACGATGAGCGAAT-3’ and 5’ - TCGGCAGGAACTTGTACTCG-3’.

### Immunostaining, Confocal Imaging, and Analysis

Immunostainings of adult brains performed as previously described (Rajan et al., 2017). Detection of InR performed as previously described (Musashe et al., 2016). Primary antibodies: chicken anti-Dilp2 (1:250; this study); rabbit anti-Dilp5 (1:500; this study); mouse anti-GFP (1:100; Sigma, cat# G6539); chicken anti-GFP (1:2000; Abcam cat#ab13970); rabbit anti-RFP (1:500; Rockland cat#600-401-379); and rabbit anti-phospho-InR β (Tyr1146) (1:1000; Cell Signaling cat#3021). Secondary antibodies: goat anti-rabbit Alexa 568 (1:500; Thermo Fisher Scientific cat#A11036); donkey anti-chicken Alexa 488 (1:500; Jackson ImmunoResearch cat#703-545-155); donkey anti-mouse Alexa 488 (1:500; Jackson ImmunoResearch cat#715-545-150); and donkey anti-chicken Alexa 647 (1:500; Jackson ImmunoResearch cat#703-605-155). Images were captured using the Zeiss LSM 800 confocal system, and analyzed with Zeiss ZenLite, ImageJ, and AIVIA (DRVISION Technologies). To measure the intensity of Dilp staining, ImageJ-calculated mean gray values from maximum intensity projections (MIPs) of a similar number of confocal stacks were averaged.

### Syt-GFP Puncta Segmentation and Analysis

Using DRVISION’s AIVIA software, we developed an in-house 3D analysis recipe to detect and segment Syt-GFP puncta. The recipe was calibrated on STAT-Gal4>UAS-Syt-GFP-expressing adult brains. To analyze the PI-STAT Syt-GFP puncta, Syt-GFP-expressing brains were co-stained with an antibody to Dilp5, and a region of interest at the point of contact between the PI-STAT neurons and the IPCs was selected from 3D projections. The segmentation recipe was run, and any Syt-GFP expression not in contact with the Dilp5-expressing processes was manually eliminated. Software analysis determined number, surface area, and volume of the segmented objects (Syt-GFP puncta). Average values were calculated for each brain, and datasets were interpreted in GraphPad. Statistical significance was quantified by 2-tailed t-test on 10-25 adult brains. Error bars represent SEM.

## Acknowledgments

We are grateful to Erika Bach, Benjamin White, Toshi Kitamoto, Kristin Scott, Ulrike Heberlein, Barret D. Pfeiffer and Richard Baines for reagents; Mary Logan for advice on InR immunohistochemistry; and Laura Holderbaum and Zach Goldberg for technical assistance. Mass spectrometry and analysis were performed by Paul Gafken in the FHCRC Proteomics Core, which is funded by an NIH Cancer Center Support Grant, P30 CA015704. Genomic reagents from the DGRC, which is funded by NIH grant 2P40OD010949, were used in this study. Stocks obtained from the Bloomington Drosophila Stock Center (NIH P40OD018537) and the Transgenic RNAi Resource project (NIGMS R01 GM084947 and NIGMS P41 GM132087) were used in this study. We thank Pin-Joe Ko for running the RNAi screen funded by the HHMI-EXORP. This study was made possible by grants awarded to AR from NIDDK (DK101605), NIGMS (GM124593), and New Development funds from Fred Hutch.

## Author Contributions

Conceptualization, A.R.; Methodology, A.R. and A.E.B.; Investigation, A.E.B.; Writing – A.R. and A.E.B.; Funding Acquisition, A.R.

---

**Figure S1.**
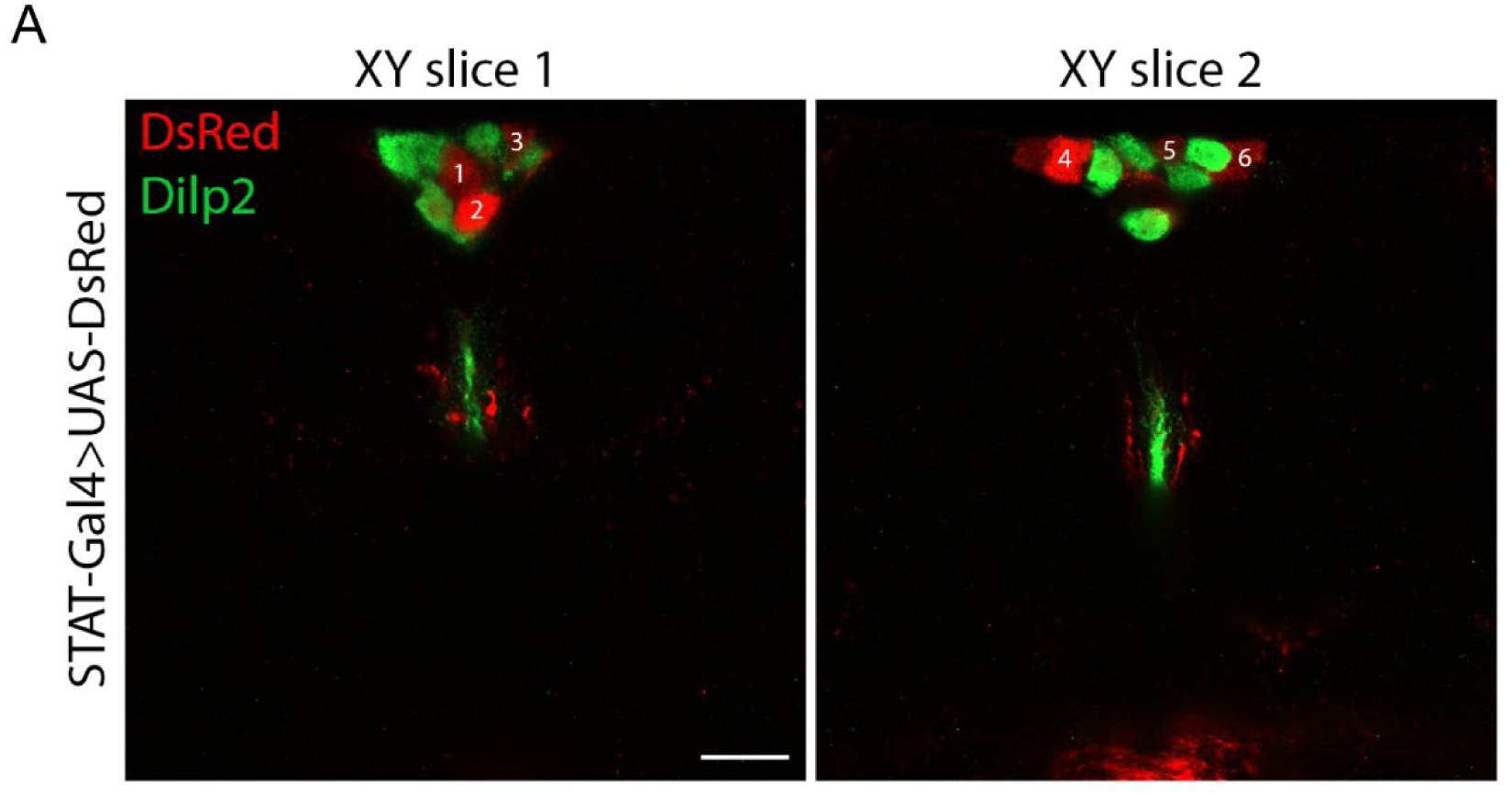
PI-STAT and IPC somas are intermingled, but not overlapping, in the PI region of the adult brain. (A) STAT-Gal4 drives expression of DsRed in a population of 6 neurons that are intermingled with the somas of the Dilp2-expressing IPCs. Scale bar, 20 μm. Dilp, *Drosophila* Insulin-like peptide.

**Figure S2.**
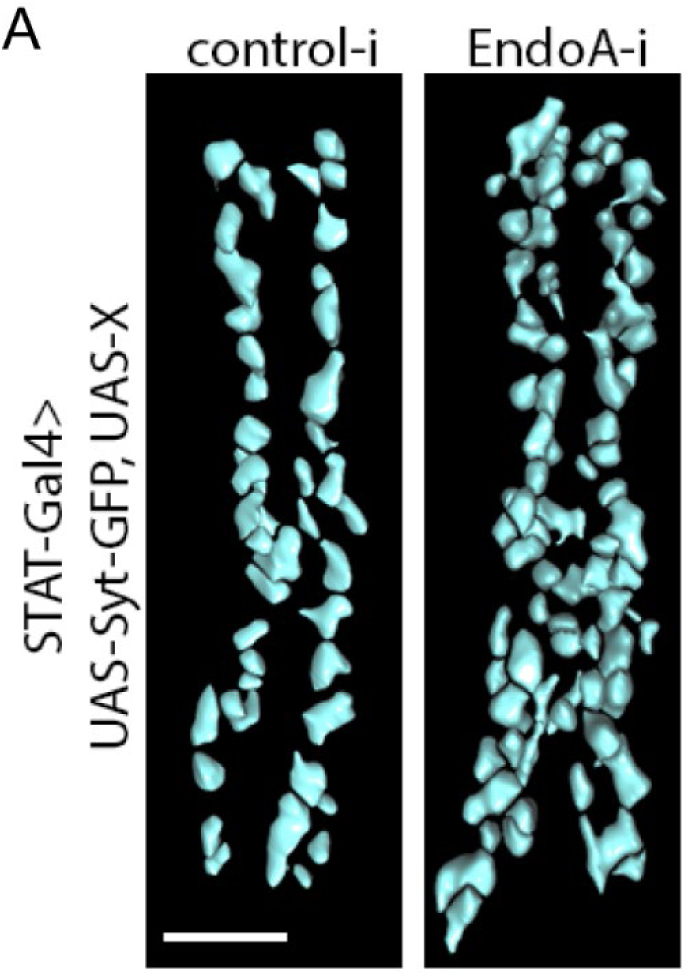
Synaptic overgrowth from knockdown of *EndophilinA* leads to increased number of Syt-GFP puncta in the PI-STAT neurons. (A) Segmentation of Syt-GFP boutons in PI-STAT neurons, following knockdown for endo via STAT-Gal4, TubGal80^ts^. Scale bar, 5 μm. PI, pars intercerebralis; Syt-GFP, synaptotagmin-GFP.

**Figure S3.**
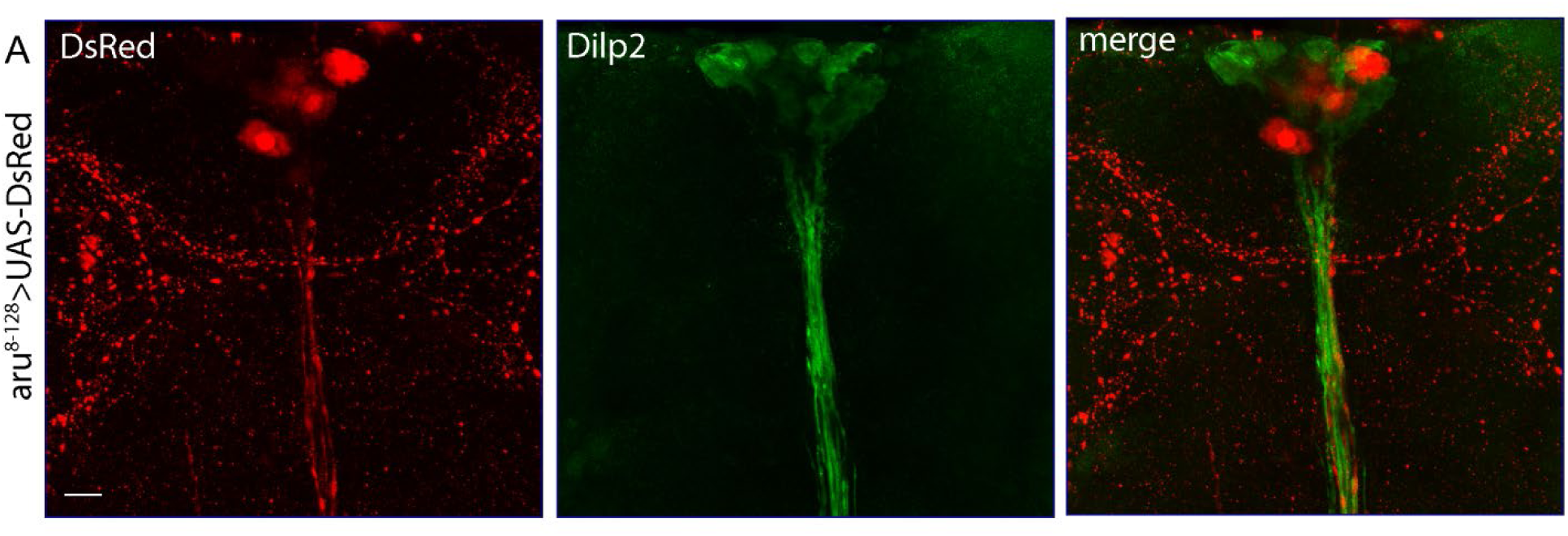
Aru is expressed in a population of PI neurons. (A) 8-128-Gal4 drives expression of dsRed in a neuron population resembling the PI-STAT neurons. Immunohistochemistry for Dilp2 identifies the IPCs. Scale bar, 20 μm. Dilp, *Drosophila* Insulin-like peptide.

**Figure S4.**
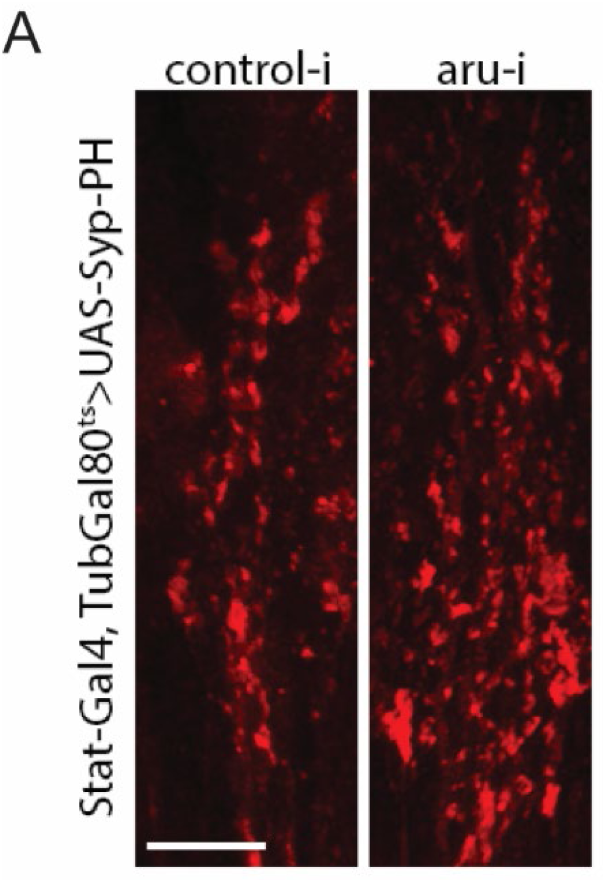
Aru regulates bouton number without affecting individual synapse activity. (A) Expression of Syp-PH in the boutons of PI-STAT neurons following control or aru knockdown via STAT-Gal4, TubGal80^ts^. *aru*-RNAi results in increased number of Syp-PH foci, but not increased expression. Scale bar, 10 μm.

**Table S1.**
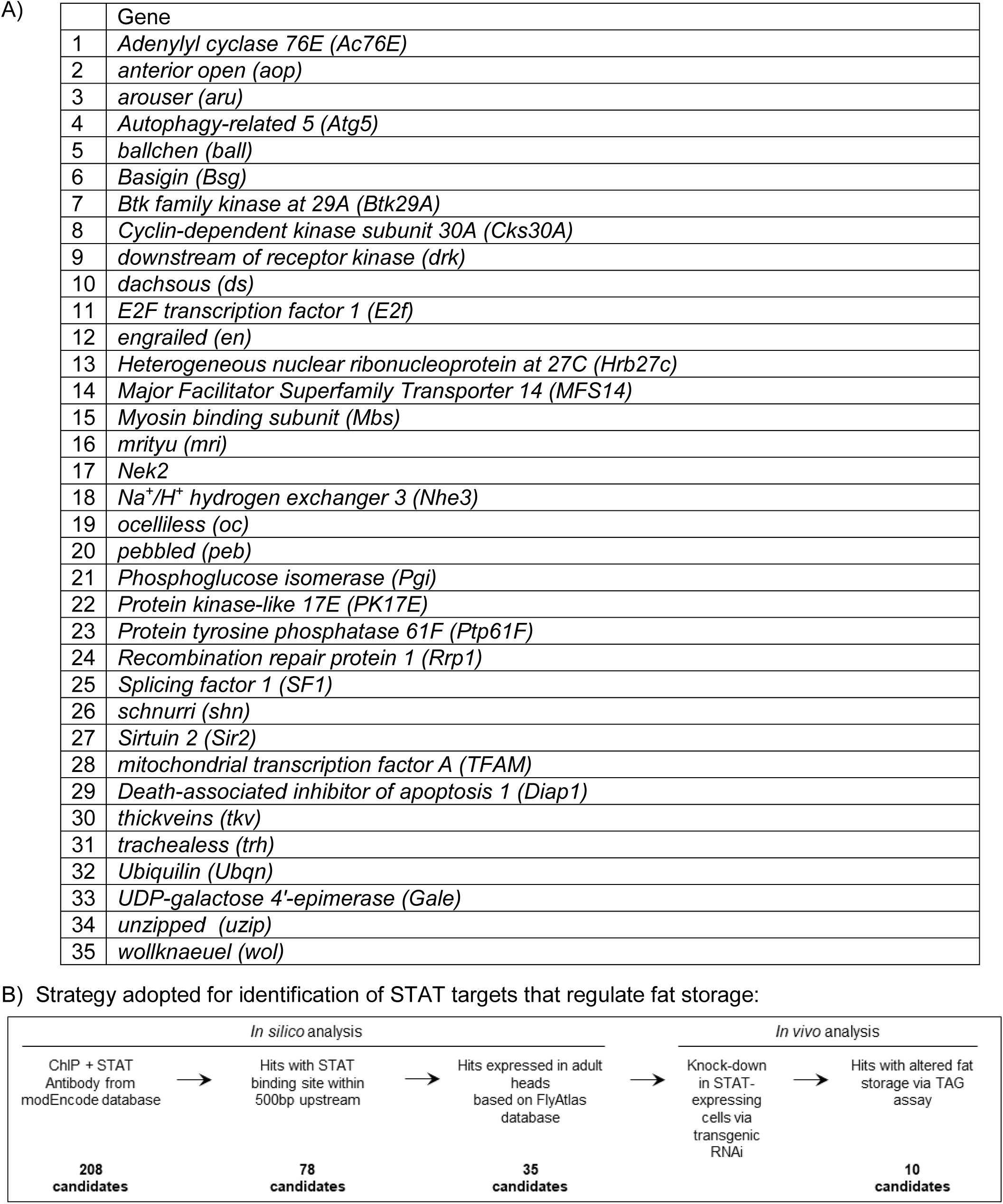
Candidate STAT target genes showing expression in the adult brain. (A) Candidate STAT targets were obtained from the modEncode database. Genes identified in FlyAtlas as being expressed in the adult brain were used for an RNAi screen. (B) Strategy adopted for identification of STAT targets that regulate fat storage. We additionally verified that our identified candidates do not affect fat storage as a result of activity within the adipocytes themselves by performing RNAi knockdown via a fat-specific Gal4 driver, and assaying for TAG levels (data not shown).

|  | Gene |
| --- | --- |
| 1 | *Adenylyl cyclase 76E (Ac76E)* |
| 2 | *anterior open (aop)* |
| 3 | *arouser (aru)* |
| 4 | *Autophagy-related 5 (Atg5)* |
| 5 | *ballchen (ball)* |
| 6 | *Basigin (Bsg)* |
| 7 | *Btk family kinase at 29A (Btk29A)* |
| 8 | *Cyclin-dependent kinase subunit 30A (Cks30A)* |
| 9 | *downstream of receptor kinase (drk)* |
| 10 | *dachsous (ds)* |
| 11 | *E2F transcription factor 1 (E2f)* |
| 12 | *engrailed (en)* |
| 13 | *Heterogeneous nuclear ribonucleoprotein at 27C (Hrb27c)* |
| 14 | *Major Facilitator Superfamily Transporter 14 (MFS14)* |
| 15 | *Myosin binding subunit (Mbs)* |
| 16 | *mrityu (mri)* |
| 17 | *Nek2* |
| 18 | *Na<sup>+</sup>/H<sup>+</sup> hydrogen exchanger 3 (Nhe3)* |
| 19 | *ocelliless (oc)* |
| 20 | *pebbled (peb)* |
| 21 | *Phosphoglucose isomerase (Pgi)* |
| 22 | *Protein kinase-like 17E (PK17E)* |
| 23 | *Protein tyrosine phosphatase 61F (Ptp61F)* |
| 24 | *Recombination repair protein 1 (Rrp1)* |
| 25 | *Splicing factor 1 (SF1)* |
| 26 | *schnurri (shn)* |
| 27 | *Sirtuin 2 (Sir2)* |
| 28 | *mitochondrial transcription factor A (TFAM)* |
| 29 | *Death-associated inhibitor of apoptosis 1 (Diap1)* |
| 30 | *thickveins (tkv)* |
| 31 | *trachealess (trh)* |
| 32 | *Ubiquilin (Ubqn)* |
| 33 | *UDP-galactose 4'-epimerase (Gale)* |
| 34 | *unzipped (uzip)* |
| 35 | *wollknaeuel (wol)* |

**Table S2.** An RNAi screen identifies candidate STAT targets that affect systemic fat storage. A table of candidate STAT targets that alter fat stores when knocked-down in STAT-expressing cells. Effect of each gene on systemic fat storage was determined via TAG assay.

| Gene | Effect on TAG | Known Functions |
| --- | --- | --- |
| *Adenylyl cyclase 76E (Ac76E)* | + | Involved in Insulin signaling; growth; response to starvation. |
| *arouser (aru)* | - | Eps8 family member; involved in response to ethanol; learning and memory; regulates synapse number. |
| *Basigin (Bsg)* | - | IgG family glycoprotein; involved in cytoskeletal rearrangements; synaptic vesicle compartmentalization and release. |
| *downstream of receptor kinase (drk)* | - | Adaptor protein; involved in Ras/Raf/MAPK signaling; cytoskeletal organization; cognition/learning. |
| *dachsous (ds)* | - | Cadherin-like; involved in Ca<sup>2+</sup>-dependent cell-cell adhesion; cell morphogenesis and differentiation. |
| *engrailed (en)* | + | Transcription factor; involved in imaginal disc development; neuroblast fate determination; axon guidance. |
| *mrityu (mri)* | - | BTB/POZ domain protein; involved in early embryo development; sensory perception of touch. |
| *Nek2* | + | Serine threonine kinase; involved in centrosome disjunction; microtubule dynamics during mitosis. |
| *pebbled (peb)* | + | Transcription factor; involved in attenuation of signal transduction; eye and tracheal development. |
| *Sirtuin 2 (Sir2)* | - | Histone deacetylase; involved in protein deacetylation; lifespan; neurodegeneration; behavioral response to ethanol. |

**Table S3.** Candidate Aru-interacting proteins identified by mass spectrometry. A table of proteins that were identified as interacting with Aru by mass spectrometry.

| Accession | Protein |
| --- | --- |
| P45888 | Actin-related protein 2 (Arp2) |
| Q9VMH2 | Actin-related protein 2/3 complex subunit 4 (Arpc4) |
| Q9VQD8 | Actin-related protein 2/3 complex subunit 5 (Arpc5) |
| O97182 | Actin-related protein 2/3 complex subunit (Arpc1) |
| P32392 | Actin-related protein 3 (Arp3) |
| Q8IQD6 | Anion exchange protein (CG8177) |
| Q8IPG9 | Basigin (Bsg) |
| E4NKG1 | Caprin, isoform B (Capr) |
| Q9VI54 | CG1965, isoform A |
| Q9V3R3 | CG3704 |
| Q8T4G5 | CG6512, isoform A |
| Q86PD2 | CG8177, isoform L |
| Q961C3 | CG8478, isoform B |
| Q59E01 | CG9795, isoform D |
| Q7KSF4 | Cheerio (Cher) |
| A1Z700 | Dilute class unconventional myosin, isoform B (Didum) |
| A0A0S0WNN6 | Embargoed, isoform B (Emb) |
| Q9VN25 | Eukaryotic translation initiation factor 3 subunit A (eIF3a) |
| B6IDY8 | Nitrogen permease regulator-like 3 (Nprl3) |
| O61604 | Fimbrin (Fim) |
| Q07171 | Gelsolin (Gel) |
| A0A0C4DHE8 | Ribosomal protein S29 (RpS29) |
| Q9W4C4 | Spoonbill (Spoon) |
| Q0KIB3 | Histone acetyltransferase type B catalytic subunit (Hat1) |
| Q9W0Q1 | Homeodomain interacting protein kinase, isoform A (Hipk) |
| Q9V3P3 | LD45860p (REG) |
| A8DYB0 | Midasin (CG13185) |
| Q9V463 | Nuclear pore complex protein Nup154 (Nup154) |
| Q9U1H9 | Nuclear RNA export factor 1 (Sbr) |
| Q9VI10 | Probable small nuclear ribonucleoprotein Sm D2 (SmD2) |
| Q7JUR6 | Protein GDAP2 homolog (CG18812) |
| Q9VN93 | Putative cysteine proteinase (CG12163) |
| A0A0B4KGG8 | Sodium/potassium-transporting ATPase subunit alpha (Atpalpha) |
| Q9VIT9 | Thioester-containing protein 4, isoform A (Tep4) |

